# PD-L1 ligation on NK cells induces a metabolic shift from glycolysis to fatty acid oxidation, enhancing tumor infiltration and control

**DOI:** 10.1101/2025.06.27.661921

**Authors:** Katja Srpan, Kyle B. Lupo, Rosa Sottile, Gianluca Scarno, Clara Lawry, Gabryelle Kolk, Brian Shaffer, Snehal G. Patel, Ahmed Al-Niaimi, Colleen M. Lau, Joseph C. Sun, Katharine C. Hsu

## Abstract

PD-L1 blockade benefits even PD-L1-negative tumors, suggesting that non-tumor cells contribute to PD-L1 expression. Natural killer (NK) cells, vital mediators of innate immunity, vigorously express PD-L1 upon activation. We demonstrate that the ligation of PD-L1 on circulating and tumor-infiltrating NK cells with the therapeutic anti-PD-L1 antibody atezolizumab, soluble PD-1, or PD-1+ cells enhances NK cell-mediated tumor clearance via changes in metabolism, adhesion, and migration. PD-L1 engagement increases NK cell tumor infiltration via the CXCR3 pathway and cytoskeletal remodeling, supported by a metabolic shift from glycolysis to fatty acid oxidation (FAO). Loss of a key FAO enzyme, CPT1A, in NK cells abrogates the PD-L1-mediated anti-tumor effect, supporting a critical role for FAO in enhanced NK cell killing. The PD-L1-triggered shift away from glycolysis permits NK cells to remain highly effective at tumor killing in glucose-restricted TME. Taken together, PD-L1 ligation enhances NK cell cytotoxicity and tumor infiltration and contributes to NK resilience in challenging TME conditions, resulting in a more effective anti-tumor immunity.

**One sentence summary:** PD-L1 engagement on NK cells enhances their tumor infiltration and cytotoxic activity by inducing a metabolic switch from glycolysis to fatty acid oxidation, enabling sustained function in the glucose-deprived tumor microenvironment.

## INTRODUCTION

The introduction of immune checkpoint inhibitors to cancer therapy has resulted in remarkable improvements in survival across various advanced tumor types ^1, 2, 3, 4, 5^. Blockade of programmed death 1 receptor (PD-1) and programmed death ligand 1 (PD-L1) stand out among immune checkpoint interventions, serving as the foundation for a growing number of immunotherapy investigations.

PD-L1 (CD274; also known as B7-H1) expression on tumor cells is induced by IFN-g secreted by activated T and NK cells ^6, 7^, creating a negative feedback mechanism to limit T cell anti-tumor activity through the induction of T cell dysfunction, exhaustion, and apoptosis ^8, 9, 10^. The prevailing paradigm within oncoimmunology, therefore, is that PD-L1 on the tumor surface effectively behaves as a molecular shield, preventing T cells in the tumor microenvironment (TME) from killing cancer cells.

While early results in melanoma patients identified PD-L1 expression on tumor cells as a predictive marker of response ^11, 12^, it was also noted that patients with PD-L1 negative tumors frequently achieved durable responses ^13, 14, 15^, leading to speculation that other cells in the TME could play a role. Correlates of response across a range of tumor types subsequently identified PD-L1 expression on tumor-infiltrating immune cells, including dendritic cells (DC), macrophages, and neutrophils, as being associated with response to anti-PD-L1 antibody treatment ^11, 13, 16^. Moreover, a study of head and neck cancer patients identified PD-L1 expression by immunohistochemistry on infiltrating immune cells as being independent of and more predictive than PD-L1 expression on tumor cells as a marker of response ^17^.

While the majority of immunotherapeutics have targeted T cells, NK cell therapies have emerged due to their versatility and lower toxicity ^18, 19, 20^. In contrast to T cells, NK cells are cytotoxic innate lymphocytes not directed by rearranged antigen-specific receptors. Instead, they distinguish stressed cells from healthy ones via a diverse repertoire of germline-encoded activating and inhibitory receptors on their surface and eradicate the aberrant cells. Despite having effector cell capacity similar to T cells, NK cells do not express high levels of PD-1 following activation ^21, 22^, but can acquire PD-1 via trogocytosis ^23^. Instead, NK cells rapidly upregulate PD-L1 upon inflammatory cytokine stimulation (IL-12, IL-15, and IL-18) or target cell activation ^24, 25^. The presence of PD-L1 on the surface of NK cells, therefore, can be considered a marker of cell activation, and the frequency of circulating PD-L1^+^ NK cells has been reported as positively associated with clinical outcomes in patients with acute myelogenous leukemia (AML) ^26^. In the same study, PD-L1^+^ NK cells exhibited more vigorous activity, and intriguingly, treatment with atezolizumab (AZ), a therapeutic anti-PD-L1 antibody that blocks PD-L1/PD-1 engagement, further increased NK cell cytolytic function. Stimulation of PD-L1 with AZ leads to NF-kB activation via p38 kinase signaling, further upregulating PD-L1 expression in a positive feedback loop ^26^.

In the present study, we show that PD-L1 is upregulated on tumor-infiltrating NK cells in patients with solid tumors. We find that engagement of PD-L1 on NK cells by AZ, soluble PD-1, or PD-1 expressed on adjacent cells leads to enhanced NK cell cytotoxicity and improved tumor clearance through surprisingly diverse mechanisms. These include priming NK cells for better migration and engagement with the tumor by enhancing cytoskeletal rearrangements and mobilizing the integrin network, increasing the surface expression of the chemokine receptor CXCR3 to augment NK cell infiltration into tumors, and inducing a metabolic shift from glycolysis to fatty acid oxidation. This metabolic versatility allows PD-L1^+^ NK cells to control the growth of highly glycolytic tumors with limited glucose availability, where T cell function is impaired.

Taken together, this work demonstrates that PD-L1 ligation not only boosts NK cell cytotoxicity but also increases tumor infiltration and improves metabolic durability in the frequently nutrient-scarce TME, ultimately resulting in enhanced anti-tumor effector function. Through this work, we offer an improved understanding of the functional role of PD-L1 in NK cells, as well as demonstrate the therapeutic potential of PD-L1 targeting in patients with PD-L1 negative tumors.

## RESULTS

### 1. NK cells in cancer patients express PD-L1

Tumor cell expression of PD-L1 and its inhibition of PD-1-expressing T cells has been extensively studied ^27^. PD-L1 expression has also been found on various immune cells, including macrophages, dendritic cells, and T cells ^28, 29, 30, 31^. Building on a previous report describing PD-L1⁺ NK cells in AML patients and their association with an activated NK cell phenotype ^26^, we identified circulating PD-L1⁺ NK cells in 62.5% of AML patients, using a threshold of >5% PD-L1⁺ NK cells to define positivity. Notably, 26% of AML patients had more than 20% PD-L1⁺ NK cells (Figure S1A). In contrast, none of the healthy donors had more than 5% PD-L1⁺ NK cells. Despite their functional similarities with T cells, circulating NK cells in the same patients did not express PD-1.

NK cells are well-recognized to be reactive to AML^32, 33, 34^, but NK cell activation in solid tumors is less well-established. To directly assess PD-L1 expression on NK cells infiltrating solid tumors, we obtained resected tumor specimens from 8 patients with oral squamous cell carcinoma (head and neck cancer, Figure 1A) and 4 with ovarian cancer (Figure 1B). While the level of tumor-infiltrating NK cells varied between tumor samples (up to 9.5% of infiltrating lymphocytes in head and neck cancer and up to 41.6% in ovarian cancer, Figure S1B), PD-L1 expression on NK cells was detected in all patients, ranging from 7 to 39% of infiltrating NK cells in head and neck cancer and from 9 to 38% in ovarian cancer (Figure 1A, B). PD-L1 was also detected on peripheral NK cells in ovarian cancer patients, but overall expression levels were lower on circulating NK cells compared to tumor-infiltrating NK cells from the same patient (Figure 1B). The expression of CD107a, a marker of NK cell degranulation, significantly correlated with PD-L1 expression on tumor-infiltrating NK cells, supporting the conclusion that PD-L1 is upregulated upon NK cell activation (Figure 1C). Expression of PD-1 on tumor-infiltrating and peripheral NK cells was minimal (Figure 1A, B).

**Figure 1:**
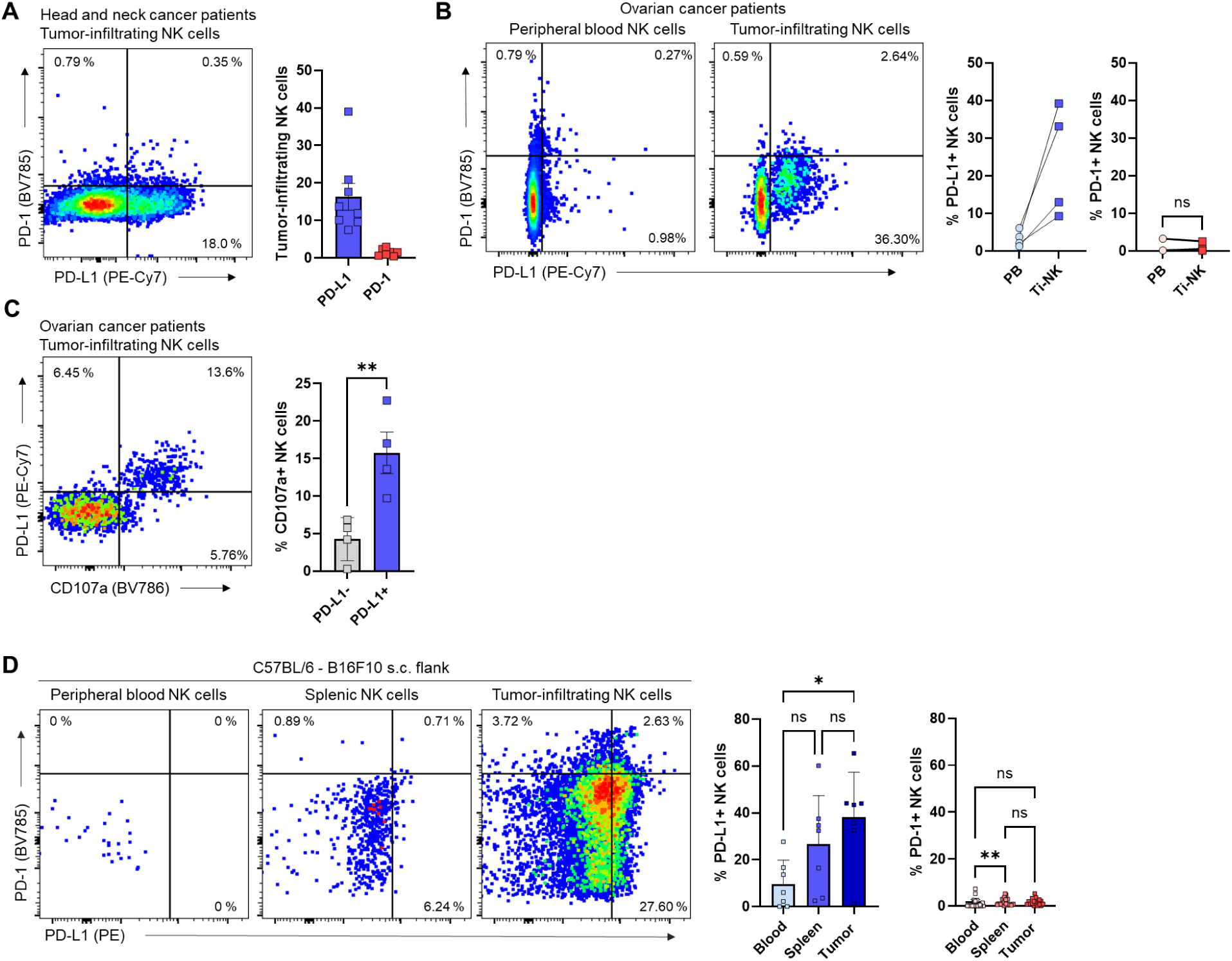
NK cells in cancer patients express PD-L1. **(A)** Representative flow cytometry plots (left) with the frequencies (right) of PD-L1^+^ and PD-1^+^ tumor-infiltrating NK cells from patients with oral squamous cell carcinoma (head and neck cancer). (n= 8 patients) **(B)** Representative flow cytometry plots (left) with the frequencies (right) of PD-L1+ and PD-1+ peripheral blood and tumor-infiltrating NK cells from patients with ovarian cancer. (n=3 patients) **(C)** Representative flow plots (left) with the frequencies (right) of CD107a^+^ PD-L1^-^ or PD-L1^+^ NK cells infiltrating ovarian tumors. (n=4) **(D)** Representative flow plots (left) with the frequencies (right) of PD-L1^+^ and PD-1^+^ circulating, splenic, and tumor-infiltrating NK cells from mice bearing B16F10 melanoma. (n=8 mice, female 4, male 4). Each symbol represents an individual patient or animal. ***P<0.001, **P<0.01, *P<0.05 with unpaired t-test (A) and Friedman test (D). Error bars, mean ± s.e.m. *See also Figure S1*.

Using a published RNA sequencing (RNA-Seq) dataset from 298 urothelial cancer tissue samples ^16^, we assessed the gene set associated with PD-L1^+^ tumor-infiltrating immune cells. While analysis of bulk transcriptomes precluded depiction of PD-L1 expression specifically on NK cells, the gene set enrichment analysis (GSEA) suggested a strong correlation between PD-L1^+^ immune cells and genes involved in cytokine, chemokine, and NK cell cytotoxicity pathways (Figure S1C, D), indicating an NK cell signature in the PD-L1^+^ immune cells.

We then corroborated the patient data using an immunocompetent mouse tumor model. Following the subcutaneous administration of B16F10 melanoma cells into the flanks of C57BL/6 mice, infiltrating immune cells were isolated and evaluated. Congruent with our findings in human primary tumor samples, we found a significant accumulation of PD-L1^+^ NK cells in tumors (Figure 1D, S1E), supporting the use of the murine tumor model to investigate PD-L1 signaling effects on NK cell function *in vivo*. PD-L1 surface expression was low but measurable on splenic NK cells and even lower on circulating NK cells. As in humans, PD-1 was not expressed by murine NK cells.

Together, these data demonstrate that PD-L1+ NK cells populate the peripheral blood in patients with hematologic malignancies and can be readily found infiltrating solid tumors. Because NK cells express PD-L1 only upon activation by target cell interaction or by cytokine induction ^26^, we conclude that the presence of PD-L1^+^ NK cells in patients with solid tumors indicates tumor-induced NK activation, either through direct cell contact or via cytokines.

### 2. PD-L1 ligation on NK cells increases their anti-tumor function

We further investigated the effect of PD-L1 ligation on NK cell-mediated control of solid tumors. We used CRISPR-Cas9 to delete *Cd274*, gene encoding PD-L1, in B16F10 melanoma (Figure S2A), thereby minimizing the possibility of tumor interaction with PD-1 on T cells. While B16F10 cells are commonly used to study NK cell activity *in vitro* and in metastatic models ^35, 36, 37^, subcutaneous solid tumor models using the same tumor cell line are NK cell-resistant ^38, 39^. We injected PD-L1^-/-^ melanoma cells subcutaneously into the flanks of C57BL/6 mice. After tumor engraftment, mice were treated with the anti-PD-L1 monoclonal antibody atezolizumab (AZ), a humanized aglycosylated IgG1 antibody that binds to both human and murine PD-L1 ^40^ or with PBS (Figure 2A). Non-treated mice grew large tumors, but AZ treatment significantly reduced tumor growth (Figure 2B, C). The depletion of NK cells completely abrogated the positive effects of AZ on tumor growth, indicating that the AZ-related anti-tumor immunity was NK cell-dependent (Figure 2C). The role of PD-L1 on NK cells was further addressed utilizing *Ncr1*Cre-*Cd274^fl/fl^* (NK-PD-L1^-/-^) mice with NK-specific *Cd274* deletion (Figure 2D). Loss of PD-L1 in NK cells did not affect the growth of the subcutaneous melanoma in untreated mice and treatment with AZ significantly suppressed tumor growth in mice with intact PD-L1 expression on NK cells. In contrast, AZ treatment of NK-PD-L1^-/-^ mice displayed tumor growth comparable to untreated controls, confirming that AZ is likely behaving in an agonistic manner to boost cytotoxicity in PD-L1^+^ NK cells. The PD-L1-mediated anti-tumor effect was further confirmed in solid PD-L1^-/-^ MC38 colon carcinoma (Figure S2B, Figure 2E, F).

**Figure 2:**
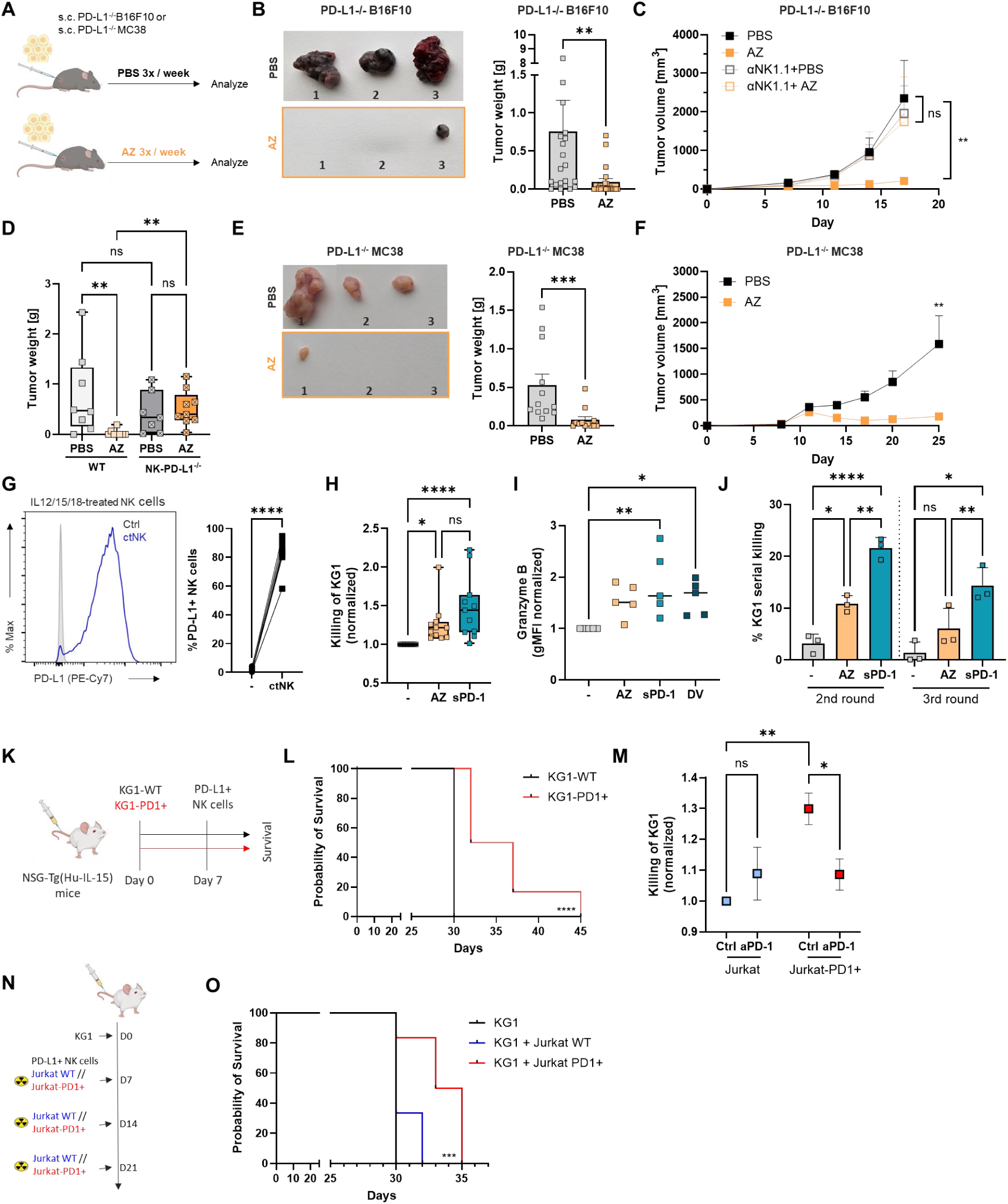
PD-L1 ligation on NK cells increases their anti-tumor function. **(A)** C57BL/6 mice were injected with PD-L1^-/-^ tumor cells (B16F10 or MC38, as indicated) subcutaneously in the right flank. Five days later, AZ was administered intraperitoneally and continued thereafter three times a week. Control mice received PBS intraperitoneally. **(B)** Representative pictures (left) and weight (right) of resected PD-L1^-/-^B16F10 tumors from mice treated with AZ or PBS. (n=20 per group, pooled from 3 independent experiments) **(C)** PBS or anti-NK1.1 treated mice were injected with PD-L1^-/-^B16F10 cells, followed by AZ or PBS administration, and tumor volume was measured at regular time points, as indicated. (n=12 per group, from 2 experiments) **(D)** *Ncr1*Cre-PD-L1 floxed mice (NK-PD-L1^-/-^) and littermate WT control were injected with PD-L1^-/-^ B16F10 and treated with PBS or AZ, as indicated in (A). Graph represents the weight of resected tumors at endpoint (n=7-9 per group) **(E)** Representative pictures (left) and weight (right) of resected PD-L1^-/-^MC38 tumors from mice treated with AZ or PBS. (n=5 per group) **(F)** C57BL/6 mice were injected with PD-L1^-/-^MC-38 cells followed by AZ or PBS administration, and tumor volume was measured at regular time points, as indicated. **(G-I)** Purified primary human NK cells were incubated with IL12/15/18 overnight (ctNK). **(G)** Representative histogram (left) and frequency of PD-L1^+^NK cells (right) in resting NK cells (Ctrl) or ctNK. (n=12 healthy human donors) **(H)** Relative killing of the KG1 cell line by ctNK or ctNK treated with AZ or soluble PD-1 (sPD-1). (n=11 healthy human donors from 4 independent experiments) **(I)** Purified primary human NK cells were incubated with IL12/15/18 (ctNK) with or without AZ or sPD-1 or durvalumab (DV) overnight. Bars represent gMFI of granzyme B (left) normalized to each donor. (n=4 healthy human donors) **(J)** Serial killing of KG1 cells by ctNK or ctNK cells treated with AZ or sPD-1. (n=3) **(K)** KG1 wild-type (WT) or KG1 transduced to express PD-1 (KG1-PD1^+^) were administered into a tail vein of NSG-Tg(Hu-IL15) mice. On day 7, mice received NK cells with constitutive expression of PD-L1 intravenously. **(L)** Survival of NSG-Tg(Hu-IL15) mice injected with WT-KG1 or PD-1^+^KG1 cells followed by administration of PD-L1^+^NK cells. (n=7 mice per condition, NK cells were isolated from 3 healthy human donors) **(M)** NK cells with constitutive expression of PD-L1 were incubated with Jurkat-WT or Jurkat-PD-1^+^ cells overnight. Where indicated, anti-PD-1 antibody nivolumab (aPD-1) was added. The following day KG1 target cells were added, and their lysis was assessed. (n=4 healthy human donors) **(N)** KG1 cells were injected intravenously into NSG-Tg(Hu-IL15) mice. On day 7, NK cells with constitutive PD-L1 expression were administered together with irradiated Jurkat-WT or Jurkat-PD-1^+^ cells via tail vein injection, followed by a weekly injection of irradiated Jurkat-WT or Jurkat-PD1^+^ cells once a week. **(O)** The survival of NSG-Tg(Hu-IL15) mice that received KG1 tumor cells and NK cells constitutively expressing PD-L1, alongside irradiated Jurkat-WT or Jurkat-PD-1^+^ cells. . (n=7 mice per condition, NK cells were isolated from 3 healthy human donors) Each symbol represents an individual mouse (B, D, E) or an individual human donor (G-J). ****P<0.0001, ***P<0.001, **P<0.01, *P<0.05 with unpaired t-test (B, E, F),Kruskal-Wallis test (D), paired t-test (G), one-way ANOVA (C), Friedman test (H, I), or two-way ANOVA (J, M), or Mantel-Cox test (L, O) . Error bars, mean ± s.e.m. (B, C, D, E, H, J, L, M) or range (D, H). *See also Figure S2*.

To assess the effect of PD-L1 ligation in human NK cells, primary NK cells were transduced to express PD-L1 constitutively (Figure S2C) and incubated with AZ or soluble PD-1 (Figure S2D, S2E). Soluble PD-1 (sPD-1) is readily detected in the media of T cells following T cell activation, as well as in the sera of patients with AML (Figure S2F). Treatment of PD-L1-expressing cells with AZ or sPD-1 led to and increased killing of the AML cell line THP-1 (Figure S2D) and significant upregulation of cytotoxic molecule granzyme B (Figure S2E). PD-L1 expression can also be rapidly induced by co-incubation with the leukemia cell line K562 ^26, 41^ or with the combination of IL12, IL15, and IL18 (cytokine-treated NK cells; ctNK), commonly used to generate the cell therapy cytokine-induced memory NK cells ^25, 42, 43^ (Figure 2G, ^26^), without affecting the expression of PD-1 (Figure S2G). Like with cells transduced to express PD-L1, engagement of PD-L1 on ctNK cells with either AZ or sPD-1 led to enhanced killing of AML target cells KG1 (Figure 2H) and greater granzyme B expression (Figure 2I). PD-L1 engagement with an alternative therapeutic antibody, durvalumab (DV), also similarly increased granzyme B production. Further, this increased cytotoxicity was not limited to a single killing but persisted through multiple consecutive rounds of target killing (Figure 2J). The increased killing was not due to antibody-dependent cytotoxicity (ADCC); AZ is non-glycosylated, making it unable to trigger ADCC ^44^, and sPD-1 lacks an Fc portion. Accordingly, the addition of an Fc-blocking antibody had no effect on the enhanced killing (Figure S2H). In contrast, deletion of the *CD274* gene in the NK cell (Figure S2I) abrogated the AZ and sPD-1-mediated effects on NK cell cytotoxicity (Figure S2J), indicating that the increased cytotoxicity was a result of signaling through PD-L1. Taken together, these data demonstrate that PD-L1 can be activated on NK cells by both a therapeutic antibody as well as by a biologically relevant source of PD-1, resulting in robust and durable anti-tumor responses.

Because PD-1 is predominantly found on the surface of cells ^45, 46^, we next ascertained whether interaction between a PD-L1^+^ NK cell with an adjacent PD-1^+^ cell can improve NK cell function, similar to AZ or sPD-1 co-incubation. We transduced KG1 leukemia cells to constitutively express surface PD-1 (Figure S2K) and intravenously injected non-transduced (WT) or PD-1^+^KG1 into immunocompromised humanized NSG-Tg(Hu-IL15) mice. After tumor engraftment, PD-L1^+^ human NK cells were administered intravenously (Figure 2K). The overall survival of mice was not affected by the expression of PD-1 on the surface of KG1 in the absence of PD-L1^+^ NK cells (Figure S2L). In contrast, PD-1^+^KG1-bearing mice receiving PD-L1^+^NK cells had significantly higher survival compared to those bearing non-transduced KG-1, supporting PD-L1-mediated enhanced anti-tumor NK cell activity (Figure 2L). To rule out the possibility that the PD-1/PDL1 interaction may have simply facilitated effector:target proximity, we also evaluated NK cell activity in the presence of PD-1^+^ T cells, pre-activated with anti-CD2-CD3-CD28-coated beads *in vitro* (Figure S2M). Coculture with PD-1^+^ T cells also increased NK cell-mediated killing of KG1 targets (Figure S2N). Further confirming the cytotoxic boost was indeed PD-L1-mediated, co-incubation of PD-L1^+^ NK cells with PD-1-transduced Jurkat cells (Figure S2O) increased NK cell cytotoxicity above baseline against KG1, an effect that disappeared when PD-1 was blocked with the anti-PD-1 antibody nivolumab (Figure 2M). In contrast, non-transduced Jurkat cells with or without nivolumab had no effect on NK cell cytotoxicity.

We then infused PD-L1^+^ NK cells together with irradiated PD-1^+^ or WT Jurkat cells into KG1-bearing NSG-Tg(Hu-IL15) mice (Figure 2N). Mice receiving PD-1^+^ Jurkat cells had significantly higher survival, which we attributed to better NK cell-mediated anti-tumor cytotoxicity (Figure 2O), as Jurkat cells alone did not affect survival (Figure S2P). These data suggest that in the TME, bidirectional communication occurs between T and NK cells via the PD-1/PD-L1 axis, whereby activated PD-1^+^ T cells may become inhibited ^24, 47^, but NK cells become vitalized for enhanced anti-tumor effector response.

### 3. PD-L1 ligation increases NK cell adhesion

We performed RNA sequencing analysis on ctNK cells with or without AZ co-stimulation and NK cells treated with IL-15 alone (Figure 3A). When performing principal component analysis, AZ-treated cells formed a distinct cluster, and the genes driving the cluster identity are associated with cytoskeletal actin filament formation [*CRIP2* ^48^], cell adhesion, migration, and interaction with the extracellular matrix (ECM) [*ITGA1*, *VCAM1*]. GSEA performed on differential gene expression following AZ treatment highlighted an upregulation of multiple integrin pathways (Figure 3B), including *ITGA1*, encoding integrin alpha-1 (CD49a), and *ITGB1*, encoding integrin beta-1, known to promote cell motility through the interaction with the ECM ^49, 50^.

**Figure 3:**
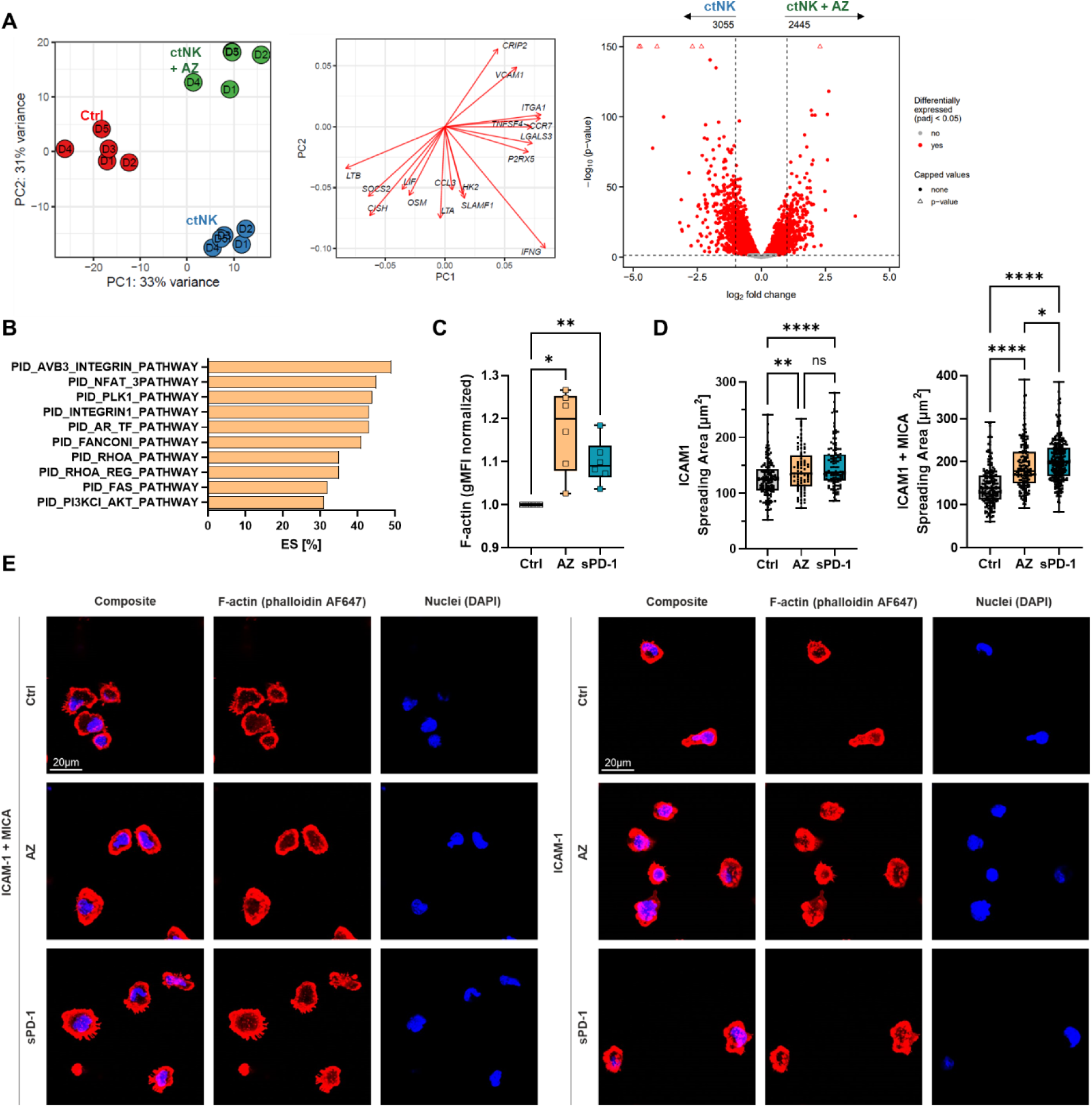
PD-L1 ligation increases NK cell adhesion. **(A)** Principal component analysis (PCA, left) and the volcano plots (right) of the RNA sequencing performed on purified human NK cells exposed to IL15 alone (Ctrl, red), IL12/15/18 (ctNK, blue) or IL12/15/18 with AZ (ctNK + AZ, green) overnight (left). PCA loading plot shows top loadings (genes) and how they contribute to each principal component (middle). (n=5 healthy human donors) **(B)** Bars plots display the enrichment score (ES) of pathways obtained from the GSEA of upregulated differentially expressed genes between ctNK and ctNK+AZ cells. **(C)** The expression of F-actin in ctNK cells (Ctrl) or ctNK cells treated with AZ or sPD-1 overnight as assessed by flow cytometry (n=6 healthy donors). **(D, E)** AZ or sPD-1-treated NK cells with constitutive PD-L1 expression and the non-treated cells were incubated for 7 mins on slides coated with ICAM-1 alone or ICAM-1 and MICA, and fixed. **(D)** The spreading area was analyzed by ImageJ. (n=3 healthy human donors). **(E)** Panels show representative confocal images of F-actin stained with phalloidin, nuclei stained with DAPI and an overlay of both. Each symbol represents an individual human donor (C) or individual cell (E). ****P<0.0001, ***P<0.001, **P<0.01, *P<0.05 with one-way ANOVA. Error bars, mean ± range. *See also Figure S3*.

Filamentous actin (F-actin) is one of the major components of the cytoskeleton responsible for cell movement, adhesion, and tumor infiltration, ^51^ and PD-L1 ligation with either AZ or sPD-1 leads to significant enhancement of the F-actin network in NK cells (Figure 3C), presumably through the upregulation of *CRIP2* (Figure S3A) ^48^. Cytoskeletal rearrangement upon engagement with a target cell has also been recognized as a critical step for effective target cell lysis ^52^, with cytotoxic lymphocytes forming a highly organized immunological synapse with an F-actin ring on its edge ^53^. The size and round shape of the immunological synapse have been associated with the strength of cytotoxic activation ^54^. We found that ctNK cells pretreated with AZ or sPD-1 formed significantly larger synapses on surfaces coated with the adhesion molecule ICAM-1 (Figure S3B, C). This was not cytokine-dependent, as AZ or sPD-1 treatment of NK cells with constitutive expression of PD-L1 also increased NK cell spreading on ICAM-1 (Figure 3D, E). This trend was maintained upon the addition of an NK cell activating ligand MICA, mimicking the surface of a target cell sensitive to NK cell activation, indicating a better overall adhesion capacity, a characteristic associated with more efficient target cell killing ^55^. PD-L1 engagement did not affect the surface expression of NKG2D, the activating receptor that binds MICA (Figure S3D), indicating that the enhanced interaction with coated surfaces was not NKG2D-mediated. Intriguingly, PD-L1 ligation did not affect the expression of CD107a, a marker of NK cell degranulation (Figure S3E), despite increasing target cell lysis (Figure 2). We conclude that PD-L1-mediated enhancement of NK cell function results from improved adhesion to target cells due to changes in F-actin rearrangement.

### 4. CXCR3-mediated recruitment to the tumor is required for enhanced tumor clearance in anti-PD-L1 treated mice

Immune cell infiltration serves as a robust positive prognostic marker for certain tumor types ^56, 57^, with NK cell infiltration, in particular, being linked to improved overall survival in solid cancers ^58, 59, 60^. One of the critical pathways for NK and T cell tumor trafficking occurs via the chemokine receptor CXCR3 ^61, 62^. The transcription of *CXCR3* gene in ctNK cells was significantly increased after treatment with AZ (Figure S4A), and both, AZ and sPD-1 increased its surface expression (Figure S4B), which also led to greater NK cell migration towards CXCL10-containing media in a trans-well experiment (Figure S4C). Treatment with AZ also induced upregulation of CXCR3 in tumor-infiltrating NK cells in mice (Figure 4A, B). The deletion of *Cxcr3* abrogated the positive effects of AZ on NK cell-mediated tumor control (Figure 4A, C, D), which was seen in the WT littermate control mice. In WT C57BL/6 mice, AZ treatment significantly increased the infiltration of NK cells into melanoma tumors, but in *Cxcr3^-/-^* mice, this increase was lost (Figure 4E). In melanoma-bearing mice that received adoptive transfer of equal numbers of NK cells from WT and *Cxcr3^-/-^* mice (Figure 4F), we observed a specific accumulation of WT NK cells in the tumor in mice treated with AZ, while the proportion of cells in the spleen was not changed (Figure 4G). Taken together, we surmise that CXCR3 contributes to the beneficial effect of AZ on NK-mediated tumor control, where higher CXCR3 expression induced by AZ leads to greater migration to the tumor and infiltration by NK cells.

**Figure 4:**
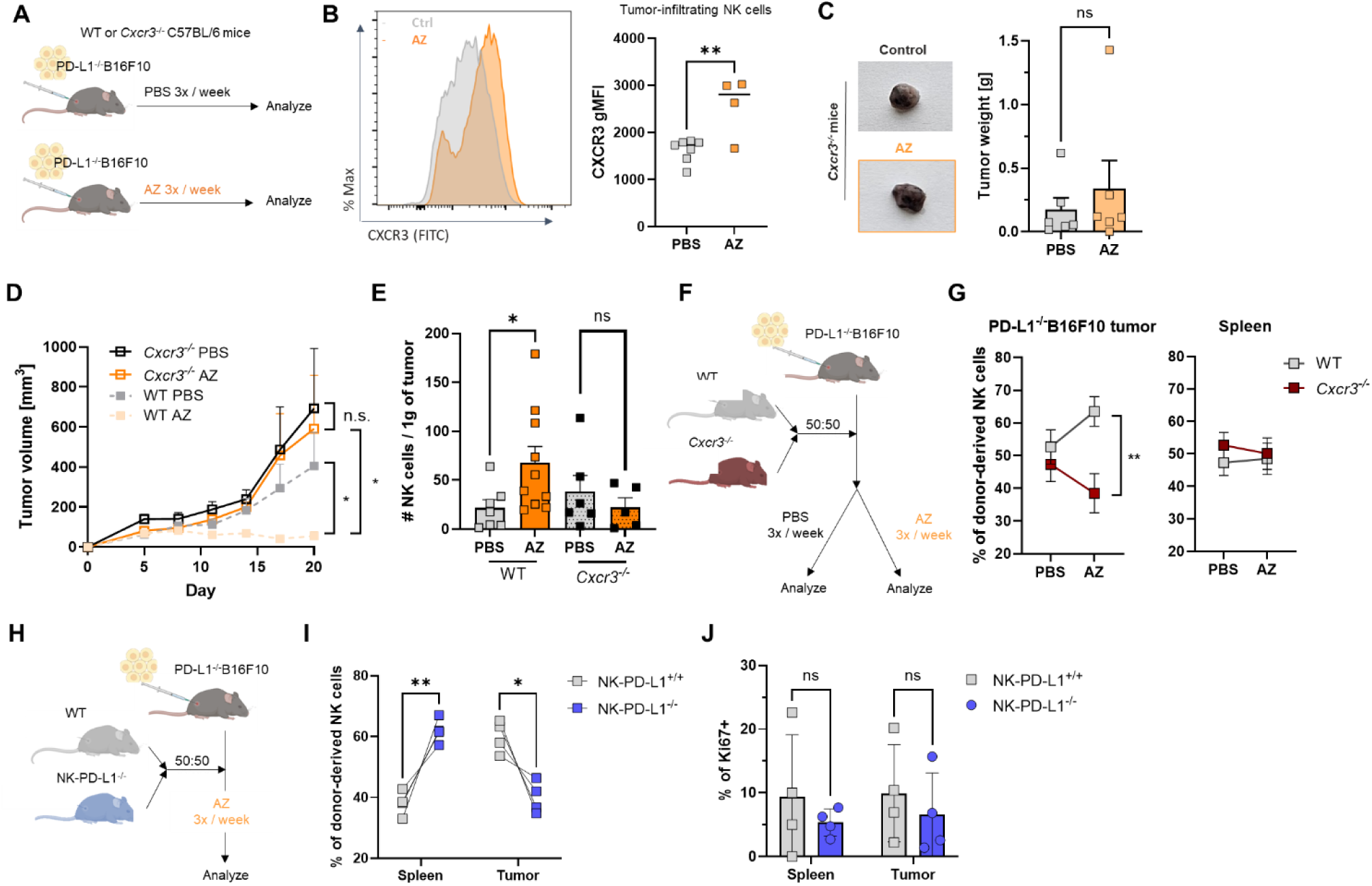
CXCR3-mediated recruitment to the tumor is required for enhanced tumor clearance in anti-PD-L1 treated mice. **(A)** *Cxcr3^-/-^* C57BL/6 mice and their WT littermates (as indicated) were injected with PD-L1^-/-^ B16F10 melanoma cells subcutaneously in the right flank. Five days later, AZ was administered intraperitoneally and continued thereafter three times a week. Control mice received PBS intraperitoneally. **(B)** Representative histogram (left) gMFI (right) of surface CXCR3 expressed by tumor-infiltrating NK cells. (n=7 per group, in AZ-treated mice, tumors from 3 mice were completely eliminated at the time of the analysis) **(C)** Representative pictures (left) and weight (right) of resected PD-L1^-/-^B16F10 tumors from mice treated with AZ or PBS. (n= 6 mice per group, pooled from 2 independent experiments) **(D)** Tumor volume was measured at regular time points, as indicated. **(E)** Tumor-infiltrating lymphocytes were isolated enumerated by flow cytometry. **(F, G)** *Cxcr3^+/+^* (*Cd45.1/.2*) and *Cxcr3^-/-^*(*Cd45.2*) NK cells were isolated and adoptively transferred into tumor-bearing CD45.1 host mice at a 1:1 ratio. Mice were then treated with AZ or PBS three times a week. **(G)** Tumor-infiltrating and splenic NK cells were enumerated 14 days post-transfer. (n=7 mice per group). **(H)** NK cells were isolated from *Ncr1Cre-CD274^fl/fl^* mice (*Cd45.2*) and their WT controls (*Cd45.1/.2*) and adoptively transferred into PD-L1^-/-^B16F10 tumor-bearing *Cd45.1* host mice at a 1:1 ratio. Mice were then treated with AZ three times a week. **(I)** Splenic and tumor-infiltrating NK cells were enumerated 7 days post-transfer. (n=4 mice per group). **(J)** The proliferation of donor-derived NK cells was assessed by Ki-67 expression. Each symbol represents an individual mouse (B, C, E, I, J). **P<0.01, *P<0.05 with non-paired t-test (B, C, D), one-way ANOVA (E) or two-way ANOVA (G, I, J). Error bars, mean ± SEM (C, D, E, G) or range (J). *See also Figure S4*.

We further validated the contribution of PD-L1 ligation to tumor homing by administering equal numbers of NK cells from PD-L1 sufficient mice and NK-PD-L1^-/-^ mice into melanoma-bearing mice (Figure 4H). Upon treatment with AZ, we observed a specific reduction of NK cells from NK-PD-L1^-/-^ mice in tumors, accompanied by an increased proportion of these cells in the spleen (Figure 4I). The proliferation of NK cells was not dependent on PD-L1 expression (Figure 4J). Therefore, PD-L1 ligation specifically impacts CXCR3-mediated NK cell tumor infiltration, the first critical component in tumor control.

### 5. PD-L1-mediated therapeutic effect requires a shift from glycolysis to fatty acid oxidation

Building on our previous identification of a link between NK cell metabolism and cytoskeletal rearrangements ^63^, we investigated whether the engagement of PD-L1 on NK cells also influences their metabolic function. Transcriptome analysis revealed that AZ-engagement of PD-L1 on ctNK significantly induced transcription of genes involved in metabolic pathways (Figure 5A, B, Figure S5A, S5B). Specifically, genes related to fatty acid oxidation (FAO) were upregulated upon AZ treatment, while genes vital to glycolysis were downregulated, indicating a metabolic shift away from glucose utilization and toward lipid metabolism. Notably, transcription of *CPT1A*, encoding a carnitine palmitoyltransferase, an enzyme critical for driving FAO, is significantly upregulated, a finding validated by increased CPT1A protein expression in ctNK following AZ and sPD-1 coincubation (Figure 5C). Treated cells also displayed higher uptake of fatty acids and lipid droplet formation (Figure 5D). Finally, mice with NK cell-specific deletion of *Cpt1a* (NK-CPT1A^-/-^) were evaluated for derangements in the clearance of PD-L1^-/-^B16F10 melanoma cells following AZ treatment (Figure 5E). While AZ treatment significantly reduced the growth of PD-L1^-/-^B16F10 melanoma of WT littermates, NK-CPT1A^-/-^ mice failed to respond to AZ (Figure 5F, G), highlighting the role of FAO in PD-L1-induced-NK-mediated tumor killing.

**Figure 5:**
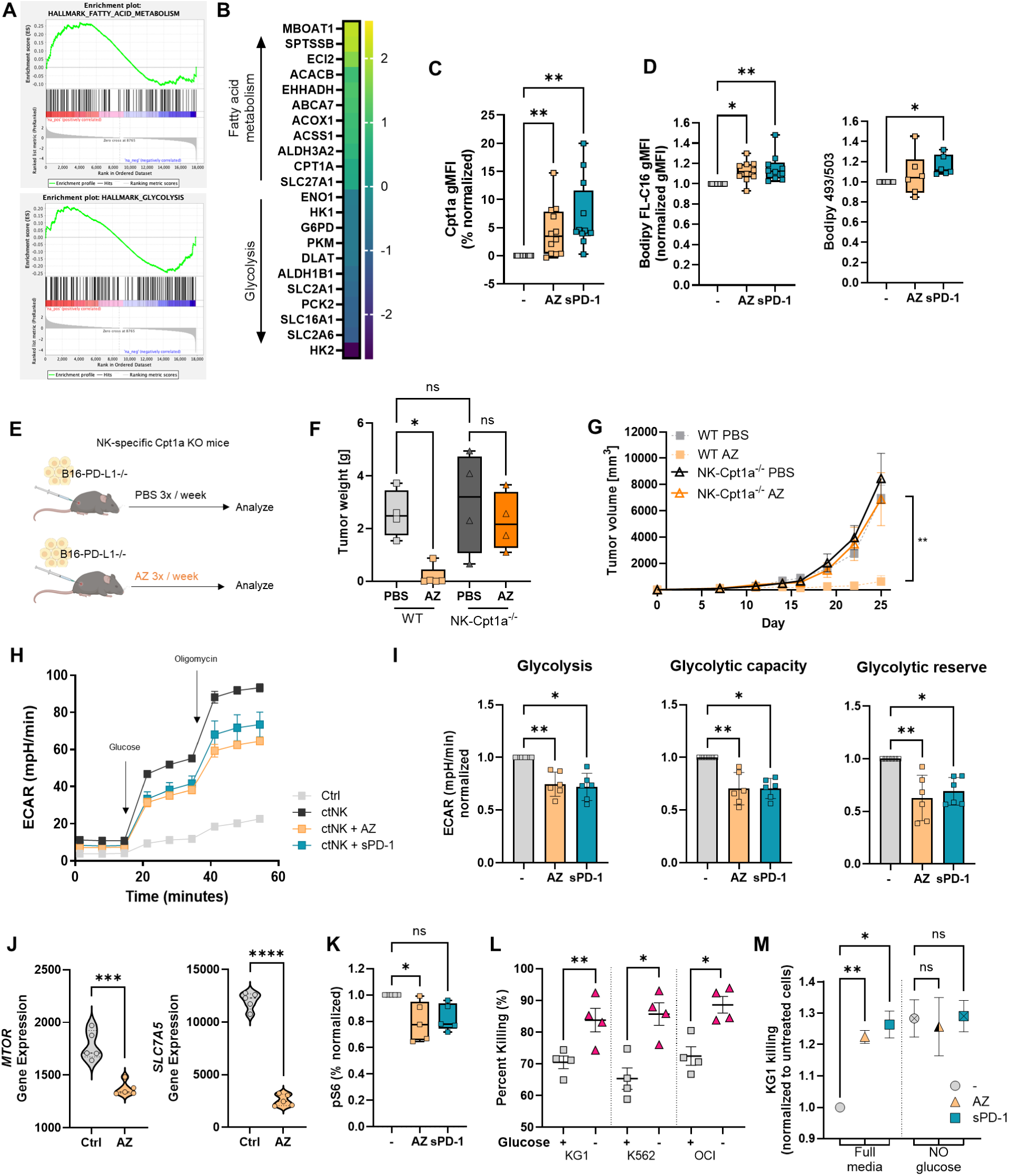
PD-L1-mediated therapeutic effect requires a shift from glycolysis to fatty acid oxidation. **(A, B)** RNA sequencing analysis was performed on ctNK cells alone or ctNK cells treated with AZ overnight. **(A)** Selected enrichment plots from pathway analysis. (n=5 healthy human donors) **(B)** Relative expression of DE genes involved in glycolysis or metabolism of fatty acids. **(C, D)** ctNK cells alone and ctNK cells treated with AZ or sPD-1 overnight were assessed for the expression of Cpt1a (n=12) **(C)** and Bodipy FLC-16 (n=12) and Bodipy 493/503 (n=6) uptake **(D)** and normalized to each donor. **(E-G)** PD-L1^-/-^B16F10 were administered into flanks of NK-specific CPT1a^-/-^ mice or their WT littermates. Mice were treated with AZ or PBS intraperitoneally three times a week. **(F)** Weight of resected PD-L1^-/-^B16F10 tumors. (n=4 mice per group) **(G)** Tumor volume was measured at regular time points, as indicated. **(H, I)** Extracellular acidification rate (ECAR) was measured in ctNK. (n=6) **(I)** Glycolysis, glycolytic capacity and glycolytic reserve were calculated from the ECAR curve and normalized to each donor. **(J)** The expression of *MTOR* (left) and *SLC7A5* (right) gene in ctNK cells and ctNK cells treated with AZ overnight. (n=5) **(K)** gMFI of pS6 in ctNK or ctNK cells treated with the AZ or sPD-1 overnight normalized to each donor. (n=6) **(L, M)** ctNK were cultured in glucose-sufficient or glucose-depleted media overnight. The lysis of indicated AML cell lines was then assessed. (n=4) **(M)** The killing capacity of ctNK or ctNK treated with AZ or sPD-1 cultured in glucose-sufficient or glucose-depleted media of KG1 target cells was assessed. (n=4). Each symbol represents an individual human donor (C, D, I-L) or individual mouse (F). ****P<0.0001, ***P<0.001, **P<0.01, *P<0.05 Friedman test (C, D, I, K), Kruskal-Wallis test (F), paired t-test (J, L), or two-way ANOVA (M). Error bars, mean ± range (C-F), or mean ± s.e.m (G-M). *See also Figure S5, S6*.

Metabolic measurements by Seahorse assay revealed that PD-L1 ligation reduced glycolysis, glycolytic capacity, and glycolytic reserve of ctNK (Figure 5H, I). Interestingly, oxidative phosphorylation (OXPHOS) in AZ-treated cells was also reduced (Figure S5C, S5D). Because mTOR is a master regulator of cell metabolism, affecting both glycolysis ^64^ and OXPHOS ^65^, we assessed mTOR activity upon the addition of AZ or sPD-1 to PD-L1^+^ NK cells. AZ treatment reduced the transcription of both the *MTOR* gene (Figure 5J) and the gene encoding the amino acid transporter *SLC7A5*, required to maintain mTOR activity ^66, 67^. PD-L1 ligation also significantly reduced the phosphorylation of S6, a key effector molecule of mTOR signaling (Figure 5K), further supporting the mechanism of metabolic change upon engagement of PD-L1 on the activated NK cell.

High glucose consumption by tumor cells restricts nutrient availability in the TME for infiltrating CD8+ T cells, leading to impaired proliferation and function of the effector cell ^68^. Because AZ treatment reduces the glycolytic capacity of NK cells while promoting their cytotoxic function, we surmised that NK cells may not be as dependent on glucose in the same way as T cells are. Indeed, ctNK cells cultured in media without glucose performed better in killing AML cell lines than those cultured in media with glucose (Figure 5L). Depletion of glucose from cultured cells also increased the expression of CPT1A, F-actin and LFA-1 adhesion molecule (Figure S6A). Overnight culture of NK cells in glucose-depleted media led to larger NK cell synapse area with thicker actin rings surfaces coated with ICAM-1 and MICA (Figure S6B, C). Under the same conditions, PD-L1 ligation with AZ or sPD-1 did not further enhance the already increased target cell killing (Figure 6M), perhaps because the absence of glucose already induced a switch to alternative nutrients ^69, 70^, boosting their cytotoxicity.

**Figure 6:**
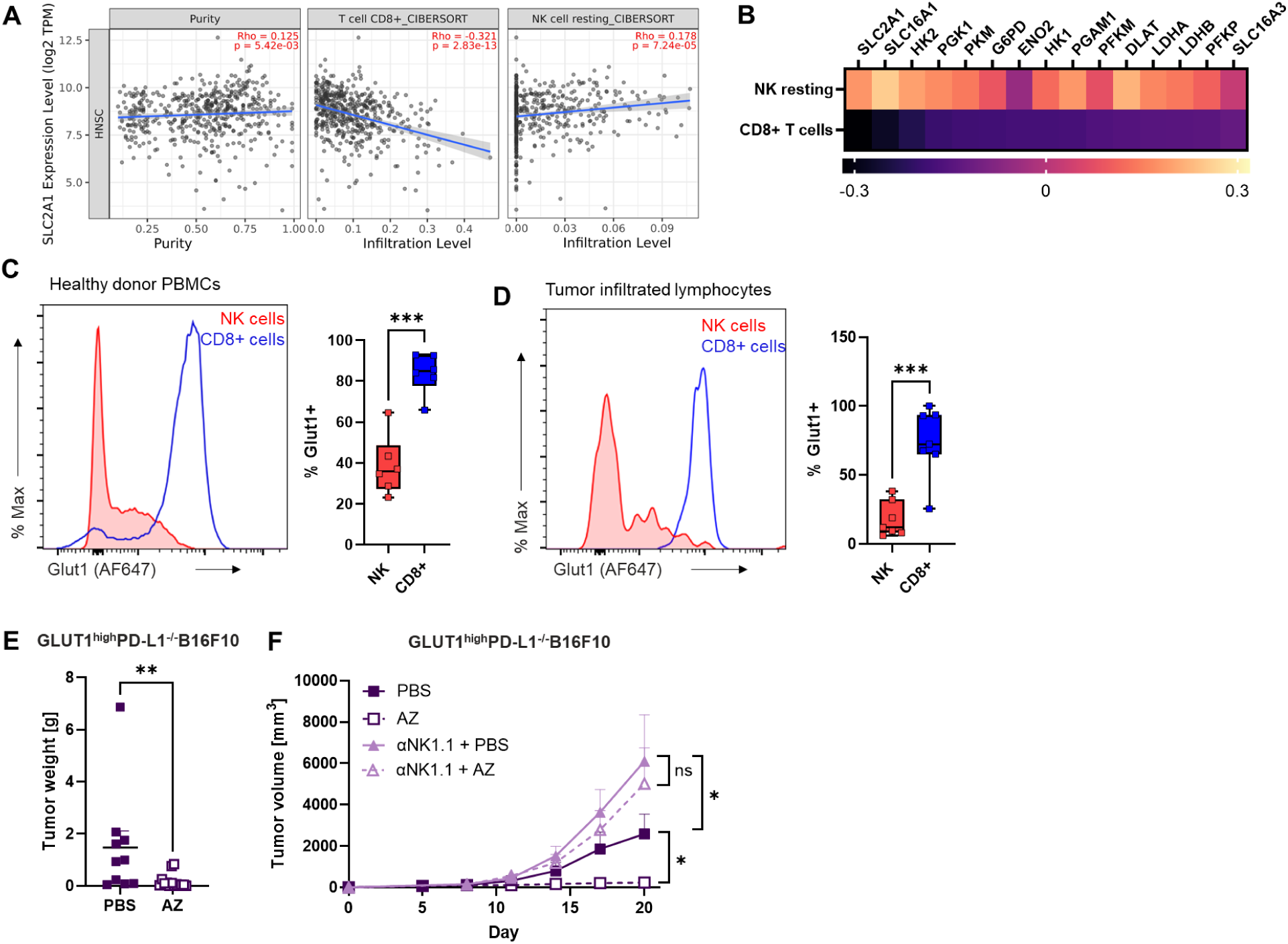
PD-L1 ligation enhances NK cell function in highly glycolytic tumors. **(A, B)** Representative plots for infiltration of CD8^+^T cells and resting NK cells relative to the expression of *SLC2A1* gene in head and neck cancer patients by Tumor Immune Estimation Resource (TIMER) web server. **(B)** Infiltration of resting NK cells and CD8^+^ T cells in head and neck tumor relative to the expression of genes involved in glycolysis. **(C, D)** Representative histogram (left) and frequency (right) of GLUT1 surface expression in circulating NK and CD8^+^ T cells from healthy donors **(C)** or NK cells and CD8^+^ T cells infiltrating head and neck tumors **(D)**. (n=6 healthy human donors and 7 head and neck cancer patients) **(E-F)** GLUT1^high^PD-L1^-/-^B16F10 were administered subcutaneously into flanks C57BL/6J mice. Mice were treated with AZ or PBS intraperitoneally three times a week. **(E)** Weight of resected tumors. (n=10 mice per group from 2 independent experiments) **(F)** Tumor volume was measured at regular time points, as indicated. (n=5 mice per group) Each symbol represents an individual human donor (C, D) or individual mouse (E). ****P<0.0001,***P<0.001, **P<0.01, *P<0.05 paired t-test (C, D) or unpaired t-test (E, F). Error bars, mean ± range (C, D) or mean ± s.e.m (E, F). *See also Figure S7*.

### 6. PD-L1 ligation enhances NK cell function in highly glycolytic tumors

Highly glycolytic tumors have been more challenging to treat with immunotherapy ^71^, in line with the hypothesis that higher glucose consumption of tumor cells negatively impacts the ability of T cells to infiltrate the tumor ^72^. Using the Tumor Immune Estimation Resource (TIMER) (https://cistrome.shinyapps.io/timer/), we found that expression of *SLC2A1,* the gene encoding the glucose receptor GLUT1, on tumor cells in head and neck cancer patients is negatively correlated with the infiltration of CD8^+^ T cells, but slightly positively correlated with NK cell infiltration (Figure 6A). Further assessment revealed the same trend for other glycolysis-related genes (Figure 6B).

We then assessed the expression of GLUT1 in circulating CD8^+^T and NK cells from healthy individuals (Figure 6C) and in cells infiltrating head and neck tumors resected from patients (Figure 6D). In both we found GLUT1 expressed at significantly higher levels in CD8^+^T cells than in NK cells, consistent with a greater dependency on glycolysis in T cells. In contrast, NK cells from expressed higher levels of CPT1A (Figure S7A, B), again highlighting how NK cells are already poised to shift toward FAO.

Because PD-L1 engagement further reduces glucose dependency in NK cells (Figure 5), we hypothesized that AZ engagement of PD-L1 boosts NK cell function even further in highly glycolytic tumors. We generated PD-L1^-/-^B16F10 melanoma cells with high expression of GLUT1 or lactate dehydrogenase A (LDHA; Figure S7C, D). Overexpression of GLUT1 or LDHA led to higher tumor utilization of glucose (Figure S7E), mimicking that of highly glycolytic tumors found in humans ^73, 74, 75, 76^. GLUT1^high^ or LDHA^high^ tumor cells were then injected subcutaneously into the flanks of C57BL/6 mice before treatment with AZ. Similar to results in metabolically unmanipulated PD-L1^-/-^B16F10 tumors (Figure 2B, C), AZ treatment led to further suppression of tumor growth in GLUT1^high^ and LDHA^high^ tumors (Figure 6E, F, Figure S8F, SG). Again, the AZ benefit was specifically attributable to NK cells, as in vivo depletion of NK cells led to the loss of the AZ benefit. Interestingly, even without AZ treatment, NK cells were empowered to control the GLUT1^high^ and LDHA^high^ tumors, indicating either increased NK vitality in the glucose-poor environment or a greater sensitivity to NK cell killing by the highly glycolytic tumors. These data provide further evidence for the potential benefit of anti-PD-L1 therapy to boost NK anti-tumor activity, even in highly glycolytic tumors where T cells are at a metabolic disadvantage.

## DISCUSSION

PD-L1 is generally considered an inhibitory ligand upregulated by tumor cells as a mechanism for inhibiting adjacent PD-1-expressing activated T cells and evading immune clearance. While it is not expressed on the surface of circulating immune cells at steady state, its expression is detected in innate and adaptive immune cells in cancer patients or under inflammatory conditions ^17, 26, 31, 77, 78^. Current anti-PD-L1-based immunotherapy mostly focuses on restoring T cell function; however, there is accumulating evidence that PD-L1 engagement also triggers functional responses in immune cells. In macrophages, PD-L1 ligation stimulates proliferation, survival, and activation, ^28^ and in dendritic cells, it promotes their migration to the draining lymph nodes during immunization or infection ^79^. We find that binding of PD-L1 on NK cells with either the therapeutic antibody or with PD-1 protein significantly augments NK cell cytotoxic capacity and anti-tumor activity through diverse mechanisms, involving cytoskeletal, migratory, and adhesion pathways and underscored by a change in metabolic programming. In sharp contrast, PD-L1 engagement in T cells suppresses their function comparable to PD-1 binding, and its genetic ablation restores antitumor response ^31^, indicating significant differences in PD-L1-mediated signaling between the innate and adaptive immune cells.

Establishing that surface expression of PD-L1 is rapidly induced upon NK activation and that PD-L1 can signal further NK activation invites the question of whether environmental sources of PD-L1 ligand are sufficient to augment NK function. PD-1 can be detected on the surface of activated T cells and on the surface of various tumors ^80^ and its solubilized form can be measured in the serum of cancer patients ^81, 82, 83^. Our data demonstrate that both membrane-bound and soluble forms of PD-1 enhance the function of PD-L1^+^ NK cells. Increased levels of sPD-1 have been reported following the initiation of cancer therapy, providing another possibility for potentiating circulating NK cells ^84, 85, 86^. PD-L1^+^ NK cells have been previously shown to inhibit PD-1^+^ T cell proliferation ^24, 47^ and impair CD8^+^ T cell-mediated response to vaccine ^78^. These studies, combined with our data, introduce the possibility that PD-L1/PD-1 interaction may trigger a bidirectional signaling dynamic, transferring the tumor-killing power from activated T cells on their way to exhaustion to neighboring activated NK cells.

PD-1 can interact with both PD-L1 and PD-L2 and interestingly, NK cell responses elicited by sPD-1 were notably stronger than those by the anti-PD-L1 therapeutic agent, which could suggest that sPD-1 is also engaging a different molecule. However, PD-L1 ligation on NK cells transduced to express PD-L1 constitutively triggered a similar increase in function with both AZ and sPD-1; and upon PD-L1 deletion, the effects were abolished, indicating PD-L1 as the sole-binding partner contributing to the observed effects. The difference in the strength of response between AZ and sPD-1 could perhaps be explained by differences in PD-L1 binding affinity, which is significantly stronger for PD-1 ^87, 88^.

It has been established that anti-PD-L1 therapy is effective even in patients with PD-L1-negative tumors ^13, 14, 15^, indicating that the benefit is occurring due to the non-tumor presentation of PD-L1. Head and neck carcinoma is known to have significant NK cell infiltration ^89^, and recent phase 2 and phase 3 clinical trials in PD-L1–low/negative recurrent or metastatic head and neck squamous cell carcinoma revealed benefit of the anti-PD-L1 monoclonal antibody durvalumab ^90, 91, 92^. Interestingly, the addition of an anti-CTLA4 antibody to the durvalumab did not have a benefit beyond the durvalumab alone, potentially indicating that unleashed T cell immunity was not the source of the anti-tumor benefit ^90^. Tumor-infiltrating NK cells could explain these findings. For AML patients who are not candidates for allogeneic hematopoietic cell transplantation, CIMNK cells represent a promising cell therapy, with 67% overall response rates ^42, 93, 94^. CIMNK cell therapy has also shown promise in pre-clinical models for the treatment of several solid tumors, including melanoma, hepatocellular carcinoma, and non-small cell lung carcinoma ^95, 96^. Cytokine treatment robustly upregulates the PD-L1 expression, and co-administration with an agonistic antibody such as AZ would represent a highly potent NK cell treatment. This combinatorial approach is now being investigated in separate phase I clinical trials for AML patients (NCT07011004) and non-small cell lung cancer patients (NCT05334329) ^96^.

PD-L1 engagement on NK cells has pleiotropic effects, collaborating for enhanced tumor control. These effects include a higher migratory capacity, an increase in solid tumor infiltration, and increased target cell synapse formation. A considerable limitation to NK cells as a cell therapy is their relative inability to infiltrate solid tumors. ECM fibers such as collagen or elastin can impede NK cell function ^39, 97, 98^, explaining why subcutaneous models of B16F10 cells rich in ECM collagen are more resistant to NK cell clearance compared to metastatic models of the same tumor ^39^. PD-L1 engagement significantly increases the number of tumor-infiltrating NK cells in otherwise NK cell-resistant subcutaneous melanoma models likely via CXCR3, a chemokine receptor essential for NK cell homing into solid tumors ^99^. Genetic deletion of *Cxcr3* in mice abrogated AZ-mediated tumor suppression, as did deletion of PD-L1 from NK cells, further illustrating that NK cell infiltration into tumors is critical to the anti-tumor function driven by PD-L1 engagement.

Once in the tumor microenvironment, PD-L1 ligation further increases the capacity of NK cells to kill tumor cells. Interestingly, the likelihood of exocytosis of lytic granules or cytokines was unchanged, suggesting that the observed increase in target cell killing must be due to greater killing efficiency by the same number of degranulating cells. Integrins in orchestration with F-actin cytoskeleton help to establish strong adhesion to the target cell while concomitantly generating costimulatory signals that boost lymphocyte activation ^100, 101^. We observed that PD-L1 binding increased multiple integrin networks and rearranged F-actin cytoskeleton into larger and thicker rings on surfaces coated with adhesion molecule or activating ligand, suggesting stronger cytotoxic cell activation ^52, 102^. A large synapse sealed with an actin ring on its edges allows for the secretion of cytolytic granules directly into the synaptic cleft without the loss of cytotoxic proteins, resulting in a more efficient killing process ^55, 103^. Nevertheless, we also observed an accumulation of granzyme B following PD-L1 engagement, suggesting that while the same number of NK cells degranulate, each lysosome might be packed with a higher density of cytolytic molecules, resulting in a more potency with each degranulation event ^104^. It is also important to note that the secretion of cytolytic granules is not the only way NK cells kill their targets; whether PD-L1 ligation affects other cytotoxic processes deserves further exploration.

Transcriptome analysis revealed an unexpected metabolic consequence following PD-L1 engagement on NK cells, subsequently confirmed by metabolic assays to be a shift away from glycolysis and toward fatty acid metabolism. Such a shift is consistent with the observation that anti-PD-L1 treatment of tumor cells also leads to reduced glycolysis via lower mTOR signaling ^105^. Apart from anaerobic glycolysis, we found oxidative phosphorylation, and mTOR-mediated signaling were also reduced in NK cells following PD-L1 binding. The high cytotoxic function of AZ- and sPD-1-treated PD-L1^+^ NK cells despite decreased mTOR activity may reflect a PD-L1-mediated optimization of the metabolic capacity of activated NK cells, perhaps to prevent their exhaustion.

Limited glucose availability in TME impairs T cell function and infiltration capacity, even when tumors are highly antigenic ^68, 71, 105, 106, 107^. In contrast, NK cells are more adaptable to nutritional challenges ^108^, preserving cytotoxic function even in the absence of glucose ^109^. Older studies on NK metabolism identified a fatty acid-rich environment and fatty acid metabolism as being suppressive to NK function ^110, 111^. In sharp contrast, we and others have shown that fatty acid oxidation boosts NK-mediated protection from viral infection and cancer via enhanced NK resilience ^63, 112, 113^. NK cells in mice exposed to a cyclic fasting diet (CFD) with a temporal reduction of blood glucose also underwent metabolic reprogramming, resulting in decreased glycolysis and increased fatty acid uptake, similar to what is observed upon PD-L1 ligation. CFD facilitated improved anti-tumor response as well as the survival of NK cells in the TME ^69^.

Unlike CD8+ T cells, NK cell infiltration in solid tumors is not impaired when tumors exhibit high glucose utilization. Using a highly glycolytic melanoma model, we demonstrate that PD-L1 binding promotes NK cell cytotoxicity against highly glycolytic tumors. Importantly, when NK cells cannot switch to FAO due to the lack of CPT1A expression, the positive effect of PD-L1 engagement is lost. We previously demonstrated that NK cells with *CPT1A* deletion were defective in forming an activating synapse on coated surfaces ^63^. This finding takes on additional significance with the discovery that PD-L1 engagement promotes a metabolic shift to CPT1A- dependent FAO. Our data strongly suggest that fatty acid metabolism is required for cytoskeletal rearrangement and that PD-L1 facilitates the process through metabolic change.

PD-L1 ligation not only enhances NK cell cytotoxic function but also increases tumor infiltration and facilitates metabolic escape, resulting in more effective anti-tumor immunity. The finding that PD-L1-binding on activated NK cells induces preferential use of fatty acid metabolism over glycolysis lends novel insight to the downstream PD-L1 signaling in NK cells and provides a mechanism by which NK cells can thrive in a highly glycolytic tumor setting. These findings underscore the potential of targeting PD-L1 to augment NK cell-mediated anti-tumor responses, either as a monotherapy or in combination with existing immunotherapies. By enhancing NK cell function and infiltration, PD-L1-directed strategies offer promising avenues for improving cancer immunotherapy outcomes.

## MATERIALS AND METHODS

### Isolation of human PBMCs, NK cells or T cells and flow cytometry

Peripheral blood was collected from healthy donors using protocols approved by the Memorial Sloan Kettering Cancer Center Institutional Review Board (nos. 06-107 and 95-054). The tumor samples were collected and processed under Biospecimen Research Protocol Institutional Review Board (nos. 06-107 and 16-1564). Donors provided informed written consent. PBMCs were isolated by Ficoll gradient purification. NK cells were purified using an NK cell isolation kit (Miltenyi Biotec) and used fresh or expanded with irradiated K562 C9 feeder cells at a ratio of 1:1 in R10 media supplemented with 200 IU/ml IL-2 (PeproTech). T cells were purified using a T cell isolation kit and cultured in R10 media supplemented with 200 IU/ml IL-2 and expanded using T Cell Activation/Expansion Kit (Miltenyi Biotec) when indicated. Tumor resections were enzymatically digested with Collagenase/Hyaluronidase Kit and supplemented with DNAse at 10ug/ml (both from StemCell) for 20 minutes at 37°C while under constant agitation. Cells were then passed through a 70um strainer, washed, and lymphocytes were purified by Percoll gradient. All samples were treated with ACK lysis buffer (0.15M NH4Cl, 0.01M KHCO3, 0.1mM NaEDTA) for 5 minutes at room temperature to lyse red blood cells.

Where indicated, primary NK cells were transduced to express PD-L1 using retroviral backbone SFG, or PD-L1 was deleted using CRISPR/Cas9 system.

Purified cells were stained for 30 minutes at 4°C with the following antibodies: PerCP anti-human CD45 (clone 2D1), BV650 anti-human CD3 (clone UCHT1), V500 anti-human CD14 (clone M5E2), PE-Cy7 anti-human PD-L1 (clone 29E.2A3), BV785 anti-human PD-1 (clone EH12.2H7), PE anti-human granzyme B (clone GB11), APC anti-human perforin (clone dG9), APC anti-human GLUT1 (clone 202915), and BV786 anti-human CD107a (clone H4A3) from BD, ECD anti-human CD56 (clone N901) from Beckman Coulter, Alexa700 anti-human CD4 (clone SK3), BV570 anti-human CD8 (clone RPA-T8) and BV421 anti-human CXCR3 (clone G025H7) from BioLegend, PE anti-human pS6 (Ser235, Ser236, clone cupk43k) from Invitrogen, and FITC anti-mouse CPT1A (clone 8F6AE9). For intracellular staining, cells were fixed and permeabilized using Cell Fixation & Cell Permeabilization Kit (Invitrogen). For the staining of phosphorylated proteins, cells were fixed and permeabilized using BD Phosflow Fix Buffer I and Perm Buffer III (BD). Cells were washed with FACS buffer after staining, resuspended in FACS buffer, and analyzed on a BD Biosciences LSR Fortessa. Flow cytometry data was analyzed using FlowJo software (BD Biosciences).

### In-vitro stimulation

Purified human NK cells were cultured at 1x10^6/ml overnight. Where indicated, recombinant human IL-12 (10ng/ml), IL-15 (50ng/ml) and IL-18 (50ng/ml), AZ (Selleck, 50ug/ml) or recombinant human PD-1 (AbCam, ab174035, 5 ug/ml) were added.

### Mice

All mice were handled in accordance with NIH and American Association of Laboratory Animal Care standards. The following mice were housed and bred on a C57BL/6J background under specific pathogen-free conditions at the Memorial Sloan Kettering Cancer (MSKCC) barrier facility: NSG-Tg(Hu-IL15) ^114^(The Jackson Laboratory), C57BL/6J (The Jackson Laboratory) were purchased and *Cxcr3*^-/-^ ^115^, C57BL/6J-*Cd45.1* (Stem) ^116^, *Ncr1iCre* ^117^, *Cpt1a^fl^*^/fl^ ^118^, *Ncr1Cre*-*Cd274^fl^*^/fl^ ^47^ were kindly provided by J. Sun, L. Lanier, J. Chun, M. Li., P. Carmeliet, and L. Brossay respectively. For all mouse experiments, 6–10-week-old age- and sex-matched littermates were used according to approved institutional protocols. Experiments were approved and authorized by the Institutional Animal Care and Use Committee (IACUC).

### Mouse tissue processing and flow cytometry staining

The mouse spleen was harvested and passed through 40um filters using flow cytometry buffer (FACS buffer; 1x D-PBS, 2% FCS). PD-L1^-/-^B16F10-tumors were resected and enzymatically digested with Collagenase/Hyaluronidase Kit and supplemented with DNAse at 10ug/ml (both from StemCell) for 20 minutes at 37°C while under constant agitation. Cells were then passed through a 70um strainer, washed, and lymphocytes were purified by Percoll gradient. Blood was collected from mice in 100 USP units/mL (BD) heparin sodium to prevent coagulation. All samples were treated with ACK lysis buffer (0.15M NH4Cl, 0.01M KHCO3, 0.1mM Na-EDTA) for 5 minutes at room temperature to lyse red blood cells. Blood and spleen samples underwent 2 rounds of lysis. Single-cell suspensions were washed and resuspended with FACS buffer containing antibodies. Cells were stained for 30 minutes at 4°C. The following antibodies were from BioLegend: BV570 anti-mouse CD45 (clone 30-F11), PerCP anti-mouse CD3 (clone 145-2C11), BV785 anti-mouse CD45.1 (clone A20), APC-Cy7 anti-mouse CD45.2 (clone 104), BV650 anti-mouse NK1.1 (clone PK136), FITC anti-mouse NKp45 (clone 29A1.4), BV421 anti-mouse CXCR3 (clone CXCR3-173), PE anti-mouse PD-L1 (clone 10F.9G2), and BV785 anti-mouse PD-1 (clone 29F.1A12). Cells were washed with FACS buffer after staining, resuspended in FACS buffer, and analyzed on a BD Biosciences LSR Fortessa. Flow cytometry data was analyzed using FlowJo software (BD Biosciences).

### Tumor Cell Lines

Jurkat, KG1 and THP-1 cell lines were purchased from ATCC. K562 C9 cells expressing mbIL21, 41BB were generated by our laboratory. When indicated, Jurkat and KG-1 cells were transduced to express PD-1 using a lentiviral vector pHR (AddGene). All human cell lines were cultured in RPMI media (MSKCC Media Core) with 10% heat-inactivated FCS (Atlanta Biologicals) and 1% Penicillin/Streptomycin (Gemini Bio).

The B16F10 cell line was purchased from ATCC. MC-38 was originally purchased from Kerafast and generously provided J. Sun. In both B16F10 and MC-38, the CD274 gene was silenced using CRISPR/Cas9, and the PD-L1 negative cells were sorted after 48hrs stimulation with murine IFN-α (50uM). When indicated, PD-L1^-/-^B16F10 cells were transduced to express high levels of GLUT1-GFP or LDHA-GFP using a lentiviral system (both Sino Biological). GLUT1^high^ or LDHA^high^ cells were sorted based on the expression of GFP. All B16F10 cell lines were cultured in DMEM media (MSKCC Media Core) with 10% heat-inactivated FCS (Atlanta Biologicals), 1% Penicillin/Streptomycin (Gemini Bio), 2mM L-glutamine (MSKCC Media Core). MC-38 cells were cultured in DMEM media (MSKCC Media Core) with 10% heat-inactivated FCS (Atlanta Biologicals), non-essential amino acids (NEAA), 1mM Na Pyruvate (MSKCC Media Core), 2mM L-Glutamine (MSKCC Media Core), 50ug/ml gentamycin (MSKCC Media Core) and 1% Penicillin/Streptomycin (Gemini Bio).

### Generation of knockout murine cell lines and primary NK cells

Deletion of PD-L1 in B16F10 and MC38 cells lines was performed via CRISPR/Cas9 genome editing using the Lipofectamine CRISPRMax Transfection Reagent (Thermo Fisher, CMAX0003), according to manufacturer recommendations. Briefly, cells were seeded at 50% confluence in one well of a 24-well plate 24 hours prior to transfection. The next day, sgRNAs (5’-UCCACCACGUACAAGUCCUU-3’, 5’-GGUCCAGCUCCCGUUCUACA-3’, 5’-UGAGCAAGUGAUUCAGUUUG-3’; Synthego) were reconstituted in Nuclease-Free Duplex Buffer (IDT, 11-01-03-01) at 0.1 nmol/uL, mixed with TrueCut Cas9 v2 protein (Thermo Fisher, A36496) and Cas9 Plus Reagent in Opti-MEM media. CRISPRMax Reagent was diluted in Opti-MEM media, mixed with the sgRNA/Cas9 solution, and incubated for 10 minutes. After 10 minutes, complex was added to cells and incubated for 3 days. Following transfection, cells were stimulated overnight with mIFNa (50uM) and sorted for PD-L1 negative cells.

Deletion of PD-L1 in primary NK cells was perfomed via CRISPR/Cas9 genome editing using the P3 Primary Cell 4D-Nucelofector X Kit S (Lonza, V4XP-3032) and Amaxa 4D nucleofector (Lonza). Briefly, primary NK cells were expanded as previously for 4-6 days. On the day of transfection, sgRNA (5’-UUGGUUGAUUUUGUUGUAUG-3’ , 5’-UCAGGCUGAGGGCUACCCCA-3’, 5’-UCUCUUGGAAUUGGUGGUGG-3’; Synthego) or non-targeting control RNA (Thermo Fisher, A35526) were reconstituted in Nuclease-Free Duplex Buffer (IDT, 11-01-03-01) at 0.1 nmol/uL, mixed 1:1 (2.5 uL sgRNA: 2.5 uL Cas 9) with TrueCut Cas9 v2 protein (Thermo Fisher, A36496) and incubated for 20 minutes to generate RNP complexes. RNP complexes were mixed with 1 million expanded NK cells, resuspended in 15 uL of P3 solution, for a final volume of 20 uL. Cells were transferred to a cuvette and nucleofected using the DN100 setting. Following transfection, cells were rested overnight, expanded as previously, and assessed for PD-L1 expression following stimulation with IL-12, IL-15, and IL-18.

### Plasmid sequences and viral production

Plasmids (1) (MG57206-ACGLN) and (2) (MG51207-ACGLN) were purchased from Sino Biological, (3) (HG10084-ACGLN) was purchased from Sino Biological and cloned into SFG backbone, and (4) was purchased from Addgene (180790).

(1) pLV-C-GFPSpark-SLC2A1: Murine lentiviral overexpression vector of SLC2A1 (GLUT1) fused to GFPSpark tag.
(2) pLV-C-GFPSpark-LDHA: Murine lentiviral overexpression vector of LDHA fused to GFPSpark tag.
(3) SFG-C-GFPSpark-PD-L1: Human retroviral overexpression vector of PD-L1 fused to GFPSpark tag.
(4) pHR-PD-1 (Y223F, Y248F)-mGFP: Human lentiviral overexpression vector of PD-1 fused to mGFP tag.

Lentiviral supernatant for (1), (2) and (4) was synthesized using a third-generation lentiviral packaging system. Each transgene was co-transfected alongside VSV-G envelope, and Gal/Pol and Rev packaging plasmids into HEK293T (ATCC, CRL-3216) packaging cells using Lipofectamine 3000 transfection reagent (Thermo Fisher, L3000001) according to manufacturer recommendations. After 24 hours and 48 hours viral supernatant was collected and used for subsequent transductions immediately or stored at -80C.

Retroviral supernatant for (3) was synthesized using Galv9 stable producer cell line. Briefly, H29 and Galv9 retroviral producer cell lines were generously provided by Dr. Anthony Daniyan. H29 cells were transfected using the ProFection Mammalian Transfection System (Promega, E1200) according to manufacturer recommendations. After 24 hours and 48 hours, viral supernatant was collected and used to transduce Galv9 stable retroviral producer cells. For retroviral production, Galv9-PDL1 cells were seeded at 50% confluence in a T150 flask and media changed after 24 hours. After 48 hours and 72 hours viral supernatant was collected and used for subsequent transductions immediately or stored at -80C.

### Retroviral and lentiviral transduction

To generate SLC2A1 (GLUT1) and LDHA over expressing cell lines, PDL1^-/-^B16F10 and PDL1^-/-^MC38 cells were seeded at 50% confluence in a 6 well plate and allowed to attach overnight. The next day, media was removed, and 2 mL of viral supernatant was added to each well. After 24 hours this process was repeated. After 48 hours, supernatant was removed, and cells were expanded and sorted for GFP+ cells.

To generate PD-1 overexpressing cell lines, KG-1 and Jurkat T cells were seeded at 1 million cells per well in a 6 well plate in 1 mL of media per well. The same day, 2 mL of viral supernatant was added to each well and cells were spinoculated at 2000xg for 1 hr. After 24 hours this process was repeated. After 48 hours, cells were centrifuged, and media replaced. Cells were expanded and sorted for GFP^+^ cells.

To generate PD-L1 overexpressing primary human NK cells, cells were isolated and expanded as described for 5 to 7 days prior to transductions. On the day of transduction, cells were seeded at 1 million cells per well in a 6 well plate in 1 mL of media per well. The same day, 2 mL of viral supernatant was added to each well and cells were spinoculated at 2000xg for 1 hr. After 24 hours this process was repeated. After 48 hours, cells were centrifuged, and media replaced. Following transduction cells were expanded for 1 to 2 weeks prior to use.

### Experimental solid tumor models and AZ treatment

Single-cell suspensions of 5 x 10^5 PD-L1^-/-^B16F10 or PD-L1^-/-^MC-38 cells were injected subcutaneously into the right flanks of indicated strains of mice. After tumor engraftment (five days), mice were treated with AZ or PBS (control), three times per week, as outlined in Figure 2A. In some experiments, mice were depleted of NK cells using a made-in-house anti-NK1.1 depletion antibody (clone PK136) or PBS (control). Tumor volume was measured twice a week.

### Competitive adoptive transfer solid tumor model

Host mice (C57BL/6J CD45.1/.2) received 3 x 10^5 PD-L1^-/-^B16F10 cells subcutaneously in their right flanks. Competitive adoptive transfer experiments were performed by injecting an equal number of magnetically purified (negative selection, Miltenyi) splenic NK cells isolated from CD45.1/.2 WT and CD45.2 CXCR3^-/-^ mice or CD45.1/.2 WT and CD45.2 NK-PD-L1-/- mice into syngeneic CD45.1 WT recipients. Each CD45.1 mouse received 10^6 cells on day 3 post-tumor transplant. Mice were then treated with AZ or PBS (control) three times per week. Tumor infiltration of each subset was assessed by flow cytometry on day 14 post-transplant for the CXCR3^-/-^ experiment or on day 7 post-transplant for the NK-PD-L1^-/-^ experiment.

### Experimental tumor xenograft models

NSG-Tg(Hu-IL15) mice were inoculated with 5 x 10^5 KG-1 or PD-1^+^KG-1 leukemia cell line intravenously. After seven days, 3 x 10^6 primary NK cells transduced to express PD-L1 from different healthy human donors were then administered. Where indicated, irradiated 2 x 10^6 Jurkat or PD-1^+^Jurkat cells were injected intravenously once per week, as indicated in Figure 2N.

### Short-term quantitative cytotoxicity assay

The short-term cytotoxicity of PD-L1-expressing cells incubated with AZ, sPD1 or PD-1^+^ cells overnight was determined by a standard luciferase-based killing assay. A total of 1 × 10^4 target tumor cells expressing GFP-luciferase were cocultured with NK cells at different E:T ratios in triplicate in white-walled 96-well plates (Corning) in a total volume of 200 μl of cell medium. To determine NK cell serial killing cytotoxic capacity, treated NK cells were first incubated with WT target cells, and fresh target cells were added every two hours. The luciferase-expressing targets were added only in the last round of the experiment. Target cells alone were plated at the same cell density to determine the maximal luciferase expression as a reference (“max signal”), and four hours later, 50 ng of d-Luciferin (Gold Biotechnology) dissolved in 50 μl of RPMI was added to each well. The emitted luminescence of each sample (“sample signal”) was detected in a Spark plate reader (Tecan) and quantified using the SparkControl software (Tecan). Percent lysis was determined as ((Target alone signal-sample signal)/(target alone signal))x100.

### In vitro cell migration assay

5x10^5 NK cells were seeded in the upper chamber of the transwell plate (pore size: 8 μm, CorningTranswell) in a serum-free media. The lower chamber was filled with medium containing CXCL10 (5 ng/ml) or control medium. Cells were incubated at 37°C, 5% CO₂ for 4 hours to allow migration. Migrated cells on the lower membrane surface were collected and counted by flow cytometry.

### Soluble PD-1 detection

Soluble PD-1 was measured in the serum of AML patients and in the supernatant from T cells, as indicated using PD-1 ELISA kit (AbCam).

### Seahorse bioanalyzer

3 × 10^5 sorted NK cells were plated in buffer-free, glucose-free media (Seahorse Biosciences/ Agilent Technologies) with glutamine (2 mM) and sodium pyruvate (0.5 mM). OCR and ECAR measurements were made under basal conditions and following the addition of glucose (10mM), oligomycin (1 μM), FCCP (1 μM), Rotenone (1 μM), and Antimycin A (1 μM) at indicated time points and recorded on a Seahorse XFe96. All compounds added are from Seahorse Biosciences/Agilent Technologies.

### Cell Counts

123count eBeads (BD Bioscience) beads were added to single-cell suspensions prior to flow cytometry. Cell numbers were enumerated according to the manufacturer’s instructions.

### Uptake of BODIPY and neutral lipid staining

PD-L1-expressing human NK cells were treated with atezolizumab or sPD-1 overnight and approximately 2 x 10^5 cells were plated and cultured with 2ng/ml BODIPY 493/503 (ThermoFisher), 10 ng/ml BODIPY FL C16 (ThermoFisher) for 30 min at 37°C in PBS, washed with FACS buffers, stained with surface antibodies, and analyzed by flow cytometry.

### Glucose detection

Cells were cultured at 2 x 10^6 cells per ml in full media for 48hrs. Glucose was measured from the supernatant according to the manufacturer’s instructions (Glucose Assay Kit, Abcam).

### F-actin imaging

For F-actin analysis, cells were allowed to settle on slides coated with NK cell ligands at 37°C for 7 minutes, fixed in 4% PFA/PBS at RT for 20 min, and permeabilized with 0.1% Triton X-100/PBS at RT for 10 min. F-actin was stained with Alexa Fluor 647–labeled phalloidin (1:200 dilution in PBS; Invitrogen) and imaged by confocal microscopy (TCS SP5) with a 63× 1.4 NA oil-immersion objective. Images were exported to ImageJ and the spreading area was measured.

### RNA-seq processing and analysis

Total RNA was extracted using Zymo Direct-zol RNA Miniprep Kit following manufacturer’s instructions (Zymo Research, Irvine, CA, USA). RNA samples were quantified using Qubit 2.0 Fluorometer (Life Technologies, Carlsbad, CA, USA) and RNA integrity was checked using Agilent TapeStation 4200 (Agilent Technologies, Palo Alto, CA, USA). RNA sequencing libraries were prepared using the NEBNext Ultra RNA Library Prep Kit for Illumina using manufacturer’s instructions (NEB, Ipswich, MA, USA). Briefly, mRNAs were initially enriched with Oligod(T) beads. Enriched mRNAs were fragmented for 15 minutes at 94 °C. First strand and second strand cDNA were subsequently synthesized. cDNA fragments were end-repaired and adenylated at 3’ends, and universal adapters were ligated to cDNA fragments, followed by index addition andb library enrichment by PCR with limited cycles. The sequencing library was validated on the Agilent TapeStation (Agilent Technologies, Palo Alto, CA, USA), and quantified by using Qubit 2.0 Fluorometer (Invitrogen, Carlsbad, CA) as well as by quantitative PCR (KAPA Biosystems, Wilmington, MA, USA).

The sequencing libraries were multiplexed and clustered onto a flowcell on the Illumina NovaSeq instrument according to the manufacturer’s instructions. The samples were sequenced using a 2x150bp Paired End (PE) configuration. Image analysis and base calling were conducted by the NovaSeq Control Software (NCS). Raw sequence data (.bcl files) generated from Illumina NovaSeq was converted into fastq files and de-multiplexed using Illumina bcl2fastq 2.20 software. One mismatch was allowed for index sequence identification.

For analysis, paired-end reads were trimmed for adaptors, and removed of low-quality reads using Trimmomatic (v0.38) ^119^. Transcript quantification was based on the hg38 UCSC Known Gene models and performed using the quasi-mapping–based mode of Salmon (v.0.13.1) ^120^, correcting for potential GC bias. Counts were summarized to the gene level using tximport (v1.10.1) ^121^. For those samples that were sequenced across two runs, summarized reads determined by tximport were summed, and the means of average transcript length offsets calculated for each run were used for downstream differential analyses executed by DESeq2 (v1.22.2) ^122^. Genes were considered DE if they showed a false discovery rate (FDR)–adjusted P < 0.05. PCA plots show all genes with average transcripts-per-million values greater than 5. Heatmaps on selected metabolism-related genes were generated using ComplexHeatmap (v1.99.7) ^123^.

## Acknowledgments

We thank members of the Hsu and Sun laboratories for comments, discussions, technical support, and experimental assistance. We are also thankful to Santosha Vardhana, Justin Cross, and the Cell Metabolism Core Facility (MSKCC), Flow Cytometry Core Facility (MSKCC), and the Molecular Cytology Core Facility (MSKCC) for their expert opinion and help with experiments and the manuscript. We thank Laurent Brossay and Peter Carmeliet for their kind donation of NK-PD-L1^-/-^ and NK-Cpt1a^-/-^ mice respectively.

## Funding

The studies were supported with the following funding: NIH P30 and HOPP research funds.

K.S. was supported by MSKCC as the First Eagle Fellow.

C.M.L. was supported by the Cancer Research Institute as a Cancer Research Institute-Carson Family Fellow.

## Author Contributions

K.S. designed and executed study experiments, performed data analysis and interpretation, and prepared the manuscript. K.L., R.S., and G.S. helped with the experimental design and performed experiments and data analysis. C.M.L. performed transcriptomic data analysis. C.L. and G.K. provided critical assistance with the experiments. S.P. and A.A.N. collected and provided patient samples. J.C.S. and B.S. provided experimental guidance and critical review. K.C.H. designed, supervised, and interpreted the study and is the corresponding author.

## Competing interests

K.C.H. is a member of the scientific advisory board for Wugen. B.S. receives research support from Genentech.

## Data and materials availability

### Lead contact

Further information and requests for resources and reagents should be directed to and will be fulfilled by the lead contact, Katharine C. Hsu.

### Materials availability

All unique/stable reagents generated in this study are available from the lead contact with a completed material transfer agreement.

### Data and code availability

The RNA-Seq data generated in this study have been deposited in the Gene Expression Omnibus under the accession number GSE273108.

Any additional information required to reanalyze the data reported in this paper is available from the lead contact upon request.

## Supplemental information

Document S1. Figures S1–S7

