## Supplemental Data for "PD-L1 ligation on NK cells induces a metabolic shift from glycolysis to fatty acid oxidation, enhancing tumor infiltration and control"

**Supplemental information**

Figure S1: NK cells in cancer patients express PD-L1, but not PD-1
Figure S2: PD-L1 ligation on NK cells increases their anti-tumor function.
Figure S3: PD-L1 ligation increases NK cell adhesion.
Figure S4: Human NK cells upregulate CXCR3 upon PD-L1 ligation.
Figure S5: PD-L1-ligation induces a metabolic shift from glycolysis to fatty acid oxidation
Figure S6: Glucose depletion positively affects NK cell cytoskeletal rearrangements.
Figure S7: PD-L1 ligation enhances NK cell function in highly glycolytic tumors.

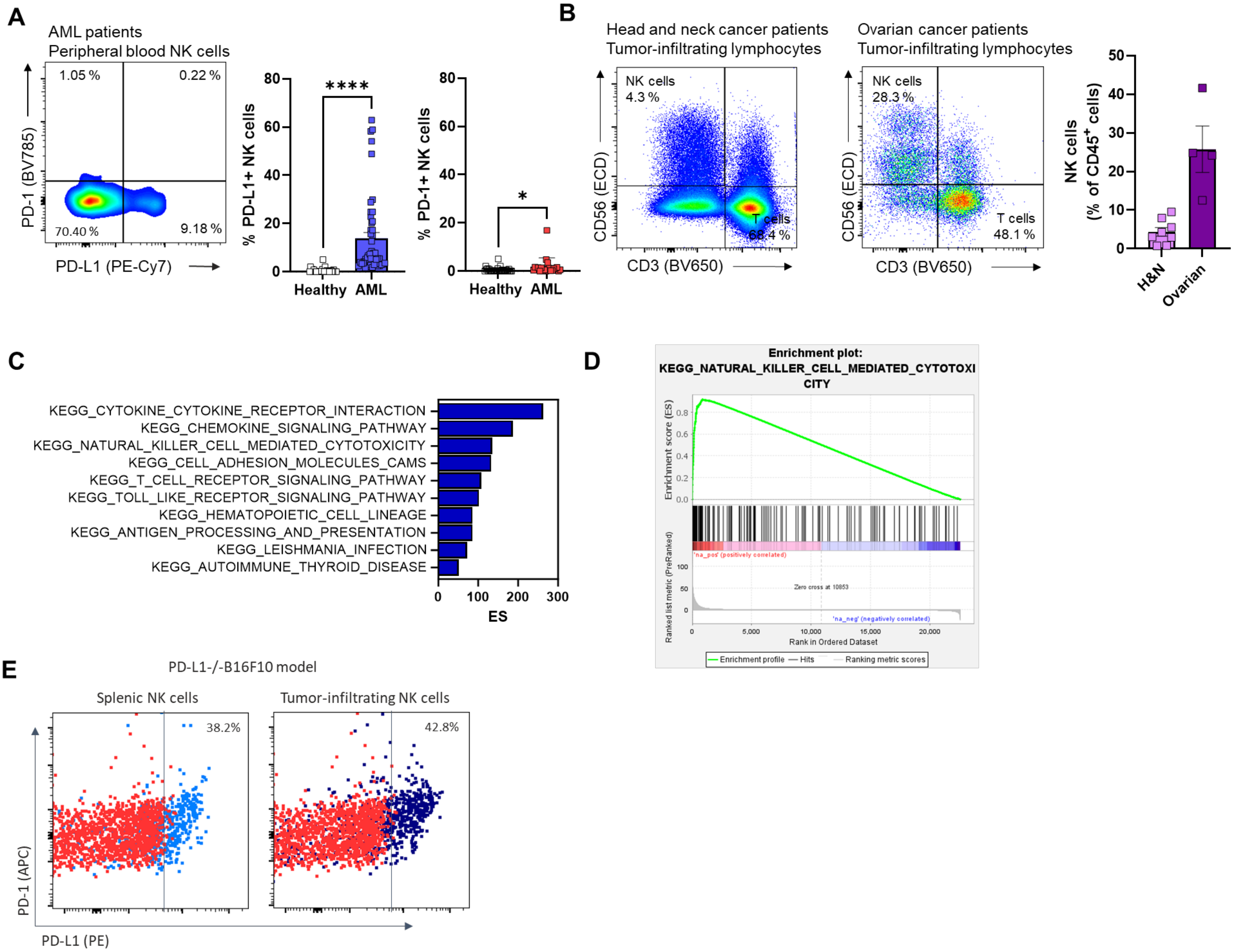


**Figure S1: NK cells in cancer patients express PD-L1, but not PD-1**

**(A)** Representative flow plot illustrating PD-L1 and PD-1 expression in peripheral blood NK cells (left) with frequencies (right) of PD-L1$^+$ and PD-1$^+$ circulating NK cells from healthy human donors or patients with AML. (n=30 healthy human donors and 48 patients with AML).

**(B)** Representative scattered plots (left) with the frequencies (right) of tumor-infiltrating lymphocytes from patients with oral squamous cell carcinoma (head and neck cancer) and ovarian cancer. (n=8 patients with head and neck cancer and 4 patients with ovarian cancer).

**(C, D)** Enriched pathways from GSEA of upregulated differentially expressed genes between PD-L1$^+$ and PD-L1$^-$ tumor-infiltrating immune cells in patients with metastatic urothelial cancer [33]. Selected GSEA plot **(D)**.

**(E)** Gating of PD-L1+ NK cells was defined based on the FMO control. The expression of PD-L1 on NK cells in splenic and tumor-infiltrating murine PD-L1$^{-/-}$B16F10 tumors. FMO control staining is depicted in red.

Each symbol represents an individual patient or human donor (A, B). ****P<0.0001, *P<0.05 with unpaired t-test (A). Error bars, mean ± s.e.m.

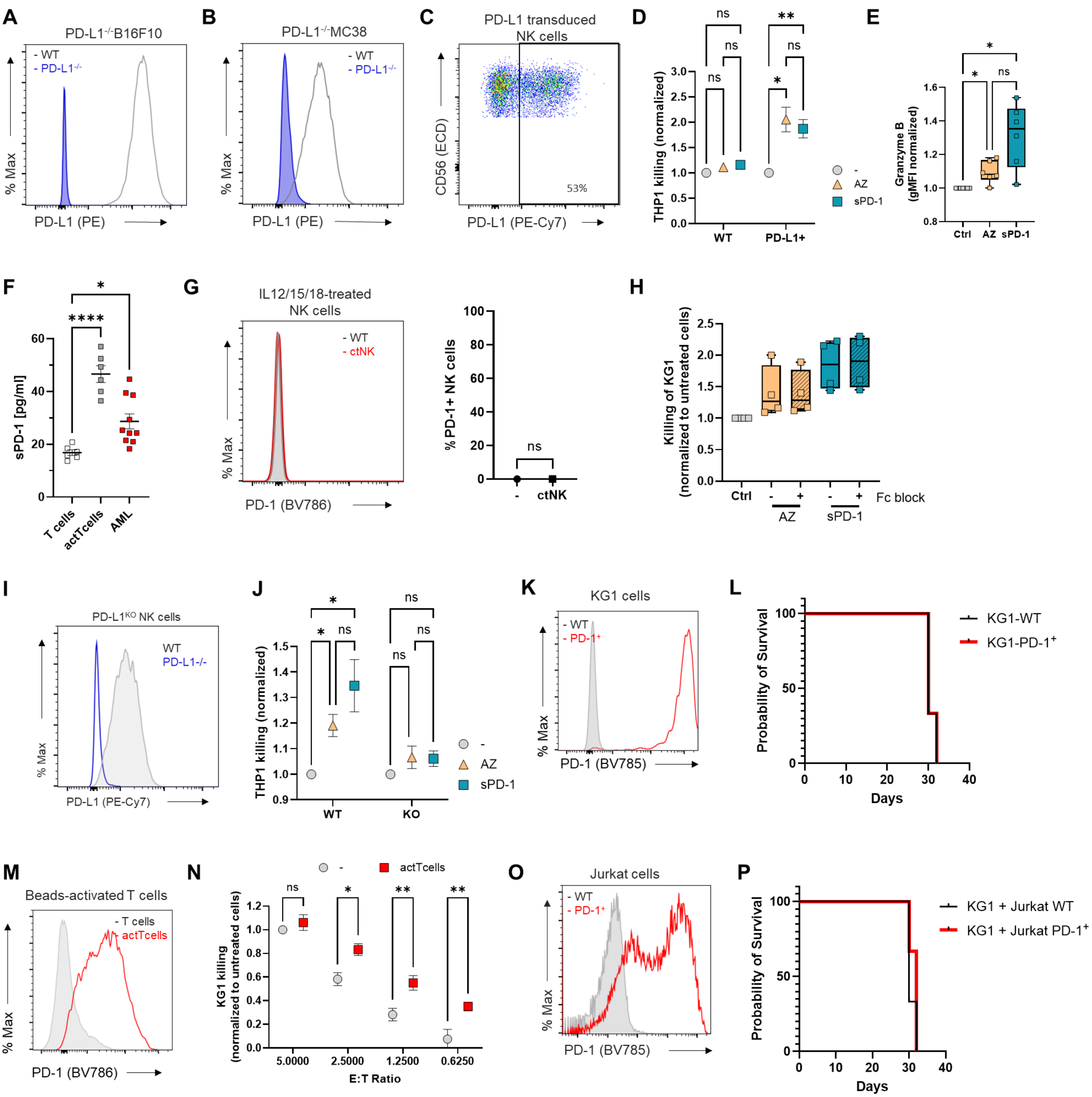
A
PD-L1-/-B16F10
- WT
- PD-L1-/-
% Max
PD-L1 (PE)
B
PD-L1-/-MC38
- WT
- PD-L1-/-
% Max
PD-L1 (PE)
C
PD-L1 transduced NK cells
CD56 (ECD)
53%
PD-L1 (PE-Cy7)
D
THP1 killing (normalized)
ns
**
ns
ns
ns
*
WT
PD-L1+
-
AZ
sPD-1
E
Granzyme B (gMFI normalized)
*
*
ns
Ctrl
AZ
sPD-1
F
sPD-1 [pg/ml]
*
****
T cells
actTcells
AML
G
IL12/15/18-treated NK cells
- WT
- ctNK
% Max
PD-1 (BV786)
% PD-1+ NK cells
ns
-
ctNK
H
Killing of KG1 (normalized to untreated cells)
Ctrl
Fc block
AZ
sPD-1
I
PD-L1KO NK cells
WT
PD-L1-/-
% Max
PD-L1 (PE-Cy7)
J
THP1 killing (normalized)
*
*
ns
ns
ns
ns
WT
KO
-
AZ
sPD-1
K
KG1 cells
- WT
- PD-1+
% Max
PD-1 (BV785)
L
Probability of Survival
KG1-WT
KG1-PD-1+
Days
M
Beads-activated T cells
- T cells
- actTcells
% Max
PD-1 (BV786)
N
-
actTcells
KG1 killing (normalized to untreated cells)
ns
*
**
**
5.0000
2.5000
1.2500
0.6250
E:T Ratio
O
Jurkat cells
- WT
- PD-1+
% Max
PD-1 (BV785)
P
Probability of Survival
KG1 + Jurkat WT
KG1 + Jurkat PD-1+
Days

**Figure S2: PD-L1 ligation on NK cells increases their anti-tumor function.**

**(A, B)** Representative histogram of PD-L1 expression upon induction with IFN-γ in PD-L1$^{-/-}$B16F10 **(A)** or PD-L1$^{-/-}$MC38 **(B)** cells or their corresponding WT controls

**(C-E)** Primary human NK cells were transduced to express PD-L1 on their surface constitutively. **(C)** Representative scattered plot of PD-L1 expression following transduction. Transduced and non-transduced NK cells were treated with AZ or sPD-1 overnight, and lysis of THP-1 **(D)** and granzyme B expression **(E)** were assessed. (n=6).

**(F)** sPD-1 concentration in the supernatant collected from non-activated T cells, T cells activated by aCD2/3/28 activation and expansion beads, or in the serum of patients with AML. (n=6 healthy donors, 10 AML patients)

**(G)** Representative histogram (left) and frequency of PD-1$^{+}$NK cells (right) in resting NK cells (Ctrl) or ctNK. (n=4 healthy human donors).

**(H)** The killing of KG1 by ctNK or ctNK cells treated with AZ or sPD-1 with or without anti-Fc antibody, as indicated, normalized to each donor. (n=4)

**(I, J)** The *CD274* gene was deleted from human primary NK cells using CRISPR/Cas9. **(I)** The histogram depicts PD-L1 expression in control vs CD274$^{-/-}$ NK cells after overnight stimulation with IL12/15/18. **(J)** Cytokine-treated CD274$^{-/-}$ cells or control NK cells were treated with AZ or sPD-1, and their killing of THP-1 cells was assessed. (n=6)

**(K)** Representative histogram of PD-1 expression in PD-1$^{+}$KG1 or WT-KG1 cells.

**(L)** The survival of NSG-Tg(Hu-IL15) mice injected with KG1- WT or KG1- PD-1$^{+}$ cells alone. (n=3 mice per group)

**(M)** Representative histogram of PD-1 expression in T cells activated by aCD2/3/28 activation and expansion beads for 3 days and non-activated T cells.

**(N)** KG1 killing at indicated effector: target ratios by CIMNK or CIMNK cells incubated with anti-CD2/3/28 bead-activated T cells (actTcells).

**(O)** Representative histogram of PD-1 expression on Jurkat cells transduced to express PD-1 (Jurkat-PD-1$^{+}$) and their WT controls (Jurkat-WT).

**(P)** The survival of NSG-Tg(Hu-IL15) mice injected with KG1 intravenously with subsequent weekly injections of irradiated Jurkat-WT or Jurkat-PD1$^{+}$ into the tail vein. (n=3 mice per group)

**P<0.01, *P<0.05 with two-way ANOVA (D, J), one-way ANOVA (N) or Friedman test (E, F). Error bars, mean ± s.e.m (D, F, J, N), or range (E, H).

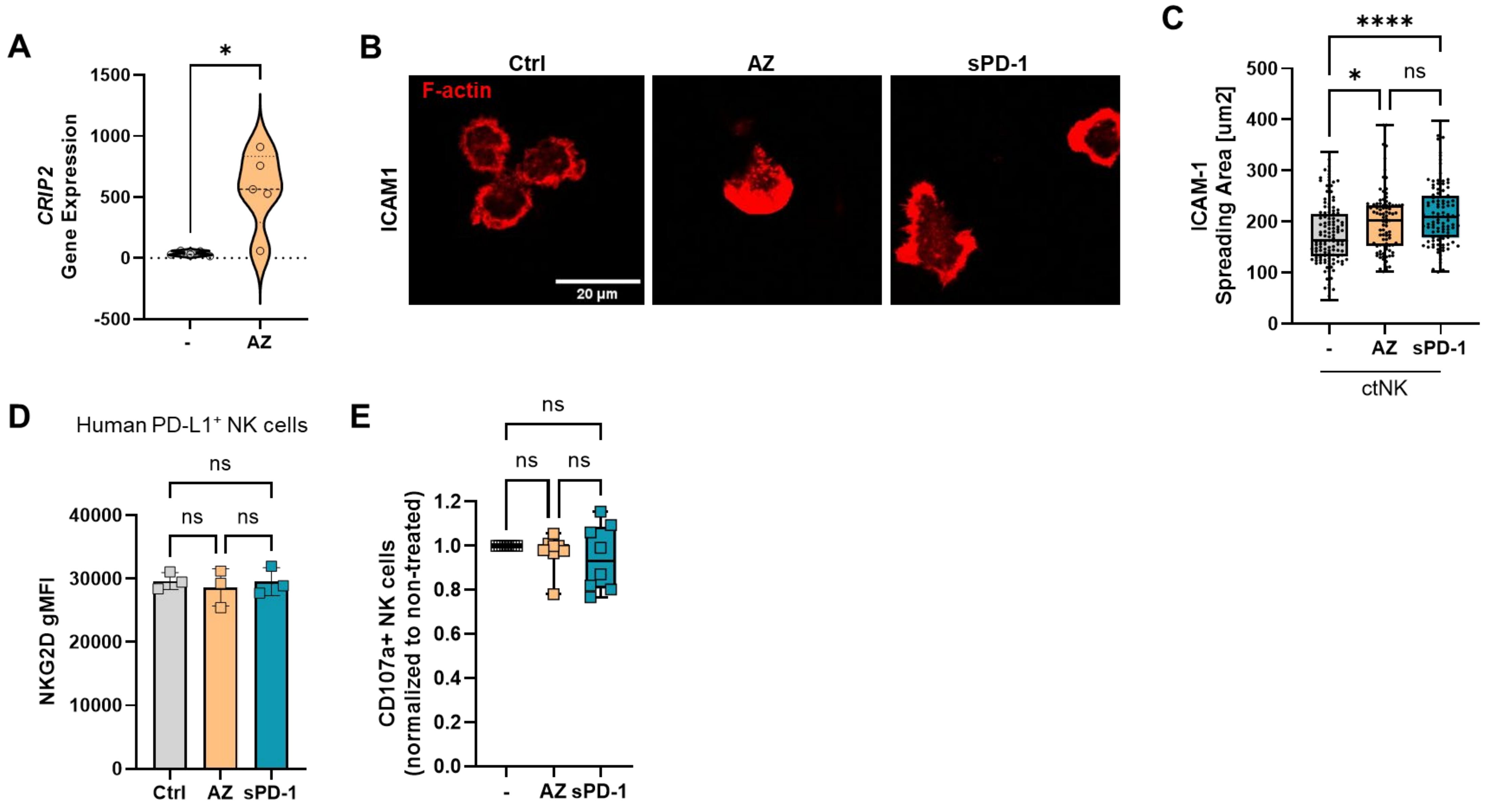


**Figure S3: PD-L1 ligation increases NK cell adhesion.**

**(A)** The expression of *CRIP2* gene in ctNK cells and ctNK cells treated with AZ overnight. (n=5).

**(B, C)** Primary human NK cells were treated with IL12/15/18 overnight alone or in presence of AZ or sPD-1-and incubated on slides coated with ICAM-1 for 7 mins and fixed. **(B)** Panels show representative confocal images of F-actin stained with phalloidin. **(C)** The spreading area was analyzed by ImageJ. (n=4 healthy human donors).

**(D)** gMFI of NKG2D in ctNK or ctNK treated with AZ or sPD-1 overnight. (n=3 healthy human donors)

**(E)** NK cells with constitutive expression of PD-L1 were treated with AZ or sPD-1 overnight. Degranulation against KG1 cells was evaluated by CD107a expression. (n=8 healthy human donors)

Each symbol represents an individual human donor (A, D, E) or an individual cell (C). ****P<0.0001, *P<0.05 with paired t-test (A), one-way ANOVA (C) or Friedman test (D, E). Error bars, mean ± range.

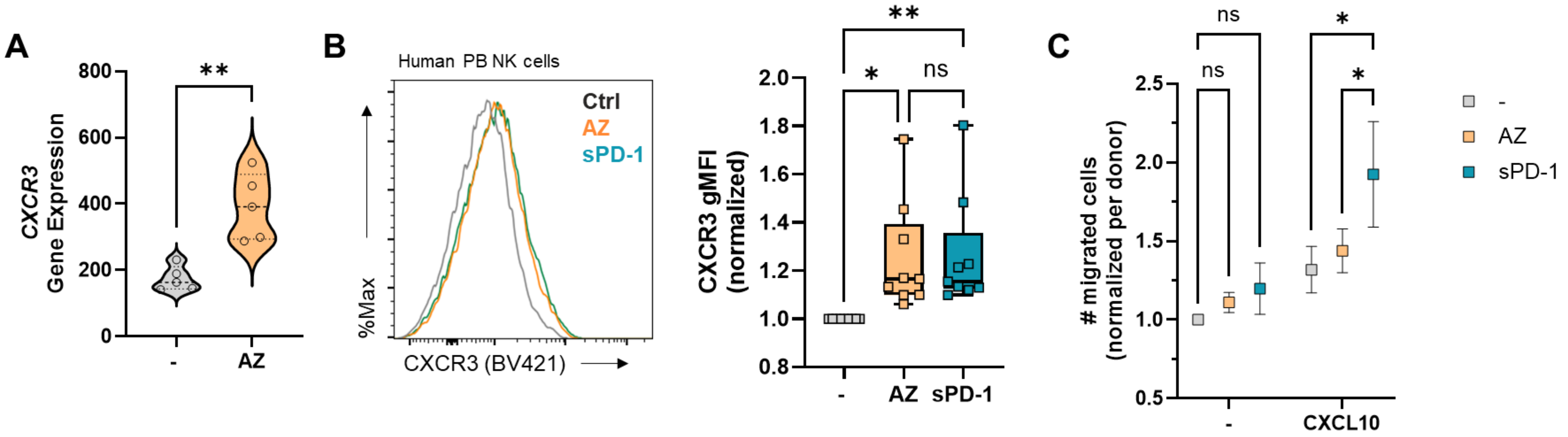


**Figure S4: Human NK cells upregulate CXCR3 upon PD-L1 ligation.**

**(A)** The expression of *CXCR3* gene in ctNK cells and ctNK cells treated with AZ overnight. (n=5).

**(B)** Representative histogram (left) and geometric mean fluorescence intensity (gMFI, right) of CXCR3 surface expression in ctNK or ctNK treated with AZ or sPD-1, as normalized to each donor. (n=9 healthy human donors)

**(C)** ctNK cells were incubated with AZ or sPD-1 overnight and transferred into a transwell plate. After 4 hours their migration into the lower compartment with or without CXCL10 was assessed by flow cytometry. (n=4 healthy human donors).

Each symbol represents an individual human donor (A, B). **P<0.01, *P<0.05 with paired t-test (A), Friedman test (B) or two-way ANOVA (C).

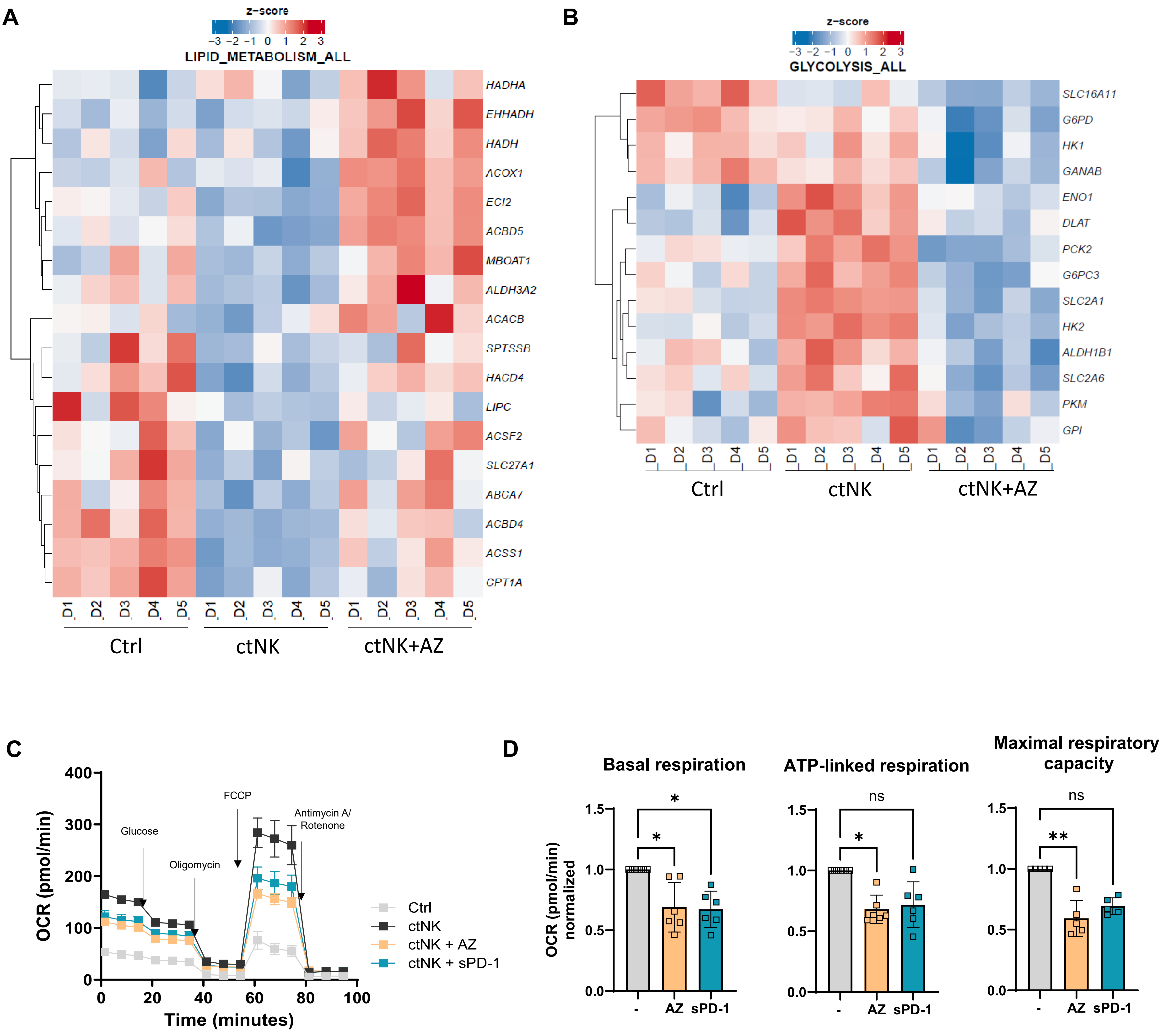


**Figure S5:. PD-L1-ligation induces a metabolic shift from glycolysis to fatty acid oxidation**

**(A, B)** Lipid metabolism **(A)** or glycolysis **(B)** related DE genes obtained from the RNA seq analysis of NK cells treated with IL-15 alone (Ctrl), IL12/15/18 (ctNK), or IL12/15/18+AZ (ctNK+AZ). (n=5 healthy human donors)

**(C, D)** Oxygen consumption rate (OCR) was measured in ctNK and ctNK cells treated with AZ or sPD-1 overnight.

**(E)** Basal respiration, ATP-linked respiration, and maximal respiration capacity were calculated from the OCR curve and normalized to each donor. (n=6 healthy human donors)

Each symbol represents an individual human donor (D) **P<0.01, *P<0.05 with Friedman test (D). Error bars, mean ± range.

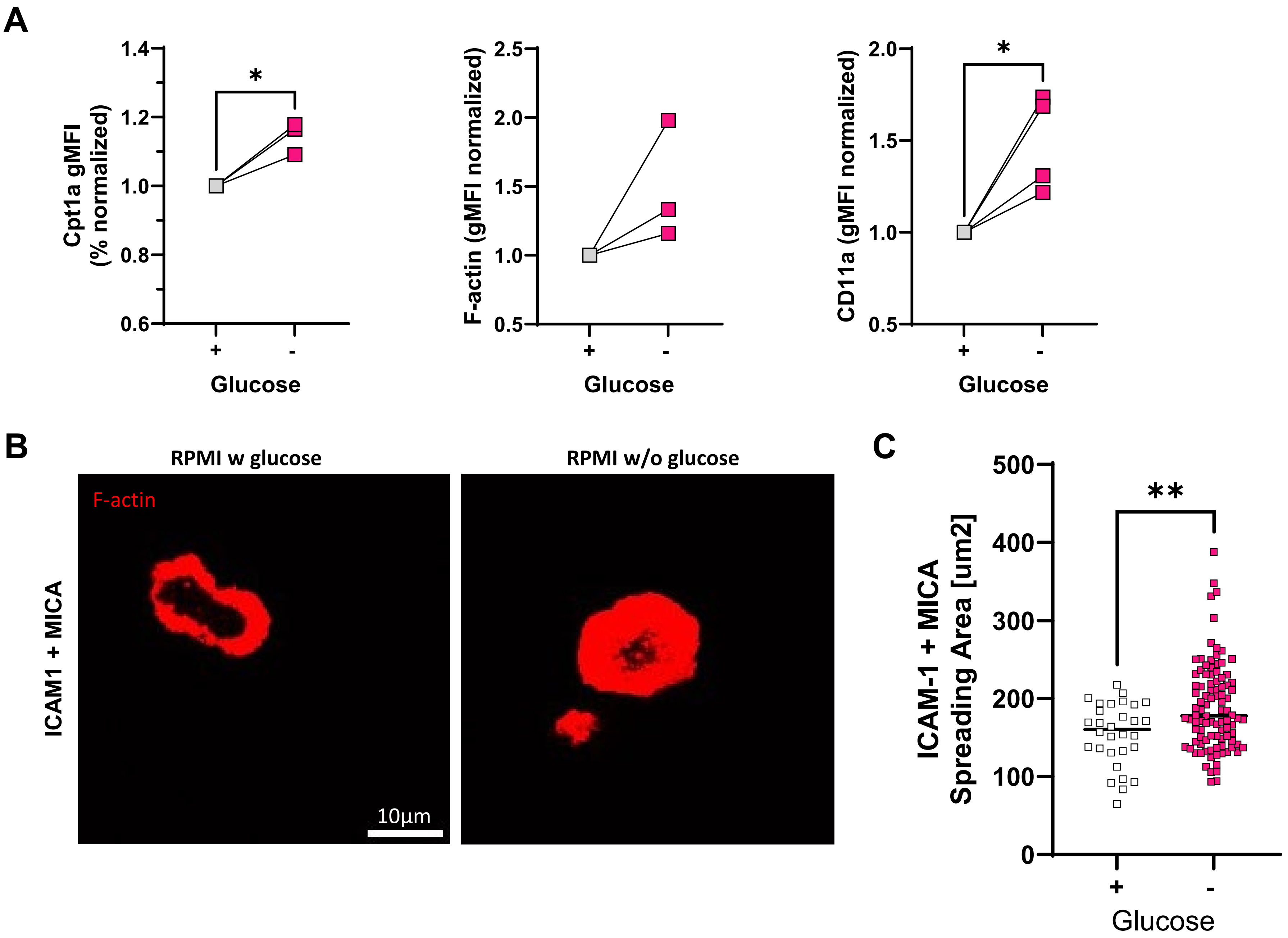


**Figure S6: Glucose depletion positively affects NK cell cytoskeletal rearrangements.**

Expanded human primary NK cells were incubated in glucose-sufficient or glucose-depleted media overnight.

**(A)** The expression of Cpt1a, F-actin and CD11a were measured by flow cytometry and normalized per each donor. (n=3)

**(B)** NK cells were placed on ICAM-1 and MICA-coated surfaces for 7 mins and fixed. F-actin cytoskeleton was stained with phalloidin and visualized by confocal microscopy. Panels are representative images from each condition.

**(C)** NK cell spreading area was measured by ImageJ. (n=3 healthy human donors).

Each symbol represents an individual human donor (A) or individual cell (C). **P<0.01, *P<0.05 with paired student t-test (A) or non-paired student t-test (C).

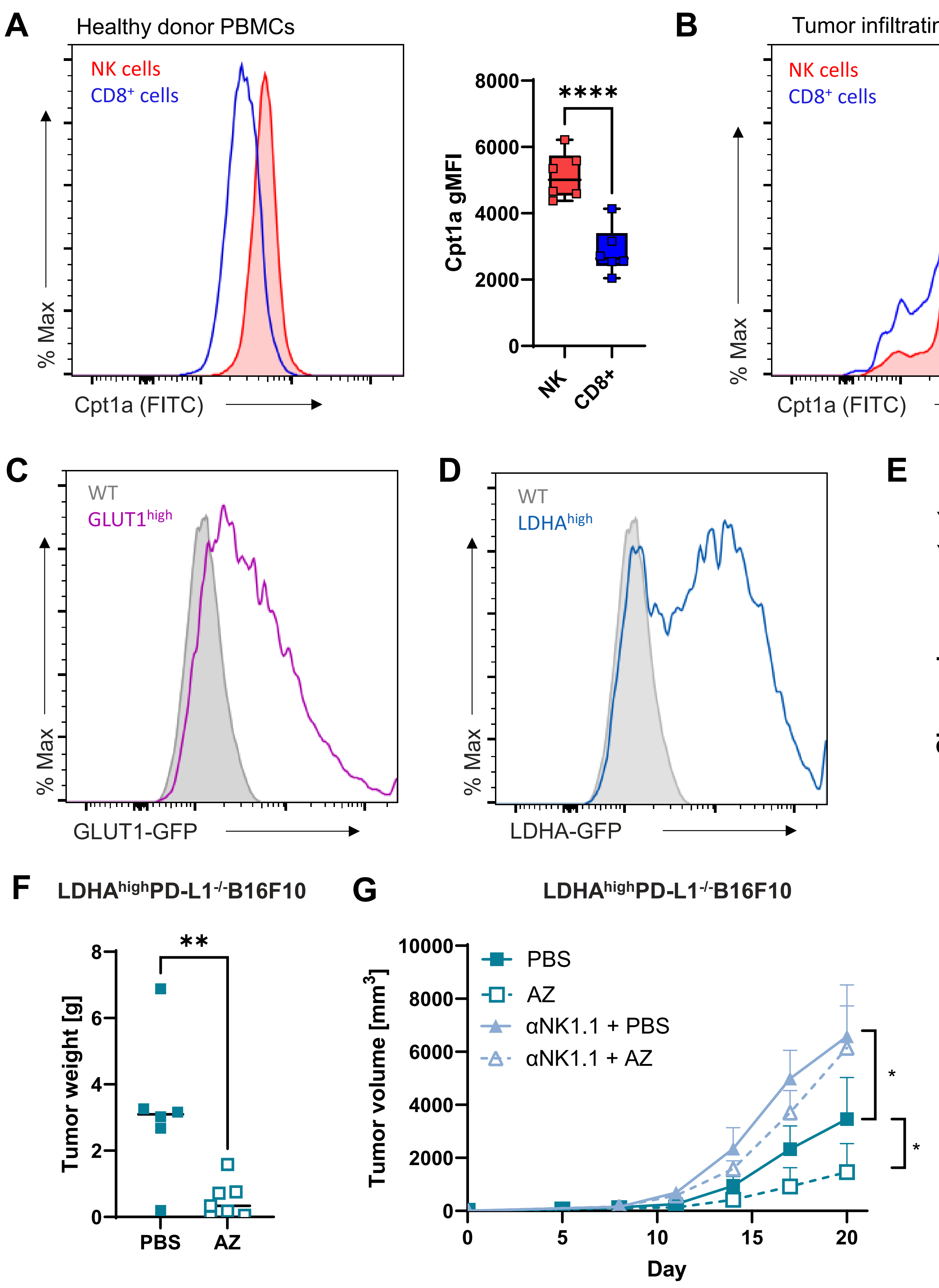


**Figure S7: PD-L1 ligation enhances NK cell function in highly glycolytic tumors.**

**(A, B)** Representative histogram (left) and gMFI (right) of Cpt1a expression in circulating NK and $CD8^+$ T cells from healthy donors **(A)** or NK cells and $CD8^+$ T cells infiltrating head and neck tumors **(B)**. (n=6 healthy human donors and 5 head and neck cancer patients)

**(C, D)** PD-L1$^{-/-}$B16F10 melanoma cells were transduced to express high levels of GLUT1 (GLUT1$^{high}$, **C**) or LDHA (LDHA$^{high}$, **D**). Representative histograms of GLUT1 **(C)** or LDHA **(D)** expression overlaid with PD-L1$^{-/-}$B16F10 control cells.

**(E)** Glut1$^{high}$ or LDHA$^{high}$ PD-L1$^{-/-}$B16F10 cells were cultured in standard media. After 48hrs the concentration of the remaining glucose was measured from the supernatant. (n=2)

**(F, G)** LDHA$^{high}$ PD-L1$^{-/-}$B16F10 were administered subcutaneously into flanks C57BL/6J mice. Mice were treated with AZ or PBS intraperitoneally three times a week. **(F)** Weight of resected tumors. (n=6 Ctrl mice and 7 AZ-treated mice) **(G)** Tumor volume was measured at regular time points, as indicated.

Each symbol represents an individual human donor (A, B) or individual mouse (F). ****P<0.0001, **P<0.01, paired t-test (A, B) and unpaired t-test (F, G). Error bars, mean ± s.e.m.
